# Cell-type and state-dependent synchronization among rodent areas S1BF, V1, perirhinal cortex and hippocampus CA1

**DOI:** 10.1101/032904

**Authors:** Martin Vinck, Jeroen J. Bos, Laura A. Van Mourik-Donga, Krista T. Oplaat, Gerbrand A. Klein, Jadin C. Jackson, Luc J. Gentet, Cyriel M.A. Pennartz

**Author notes:** Current address: Department of Neurobiology, Yale University, New Haven, USA. These authors contributed equally.

## Abstract

Beta and gamma rhythms have been hypothesized to be involved in global and local coordination of neuronal activity, respectively. Here, we investigated how cells in rodent area S1BF are entrained by rhythmic fluctuations at various frequencies within the local area and in connected areas, and how this depends on behavioral state and cell type. We performed simultaneous extracellular field and unit recordings in four connected areas of the freely moving rat (S1BF, V1M, perirhinal cortex, CA1). S1BF spiking activity was strongly entrained by both beta and gamma S1BF oscillations, which were associated with deactivations and activations, respectively. We identified multiple classes of fast spiking and excitatory cells in S1BF, which showed prominent differences in rhythmic entrainment and in the extent to which phase locking was modulated by behavioral state. Using an additional dataset acquired by whole-cell recordings in head-fixed mice, these cell classes could be compared with identified phenotypes showing gamma rhythmicity in their membrane potential. We next examined how S1BF cells were entrained by rhythmic fluctuations in connected brain areas. Gamma-synchronization was detected in all four areas, however we did not detect significant gamma coherence among these areas. Instead, we only found long-range coherence in the theta-beta range among these areas. In contrast to local S1BF synchronization, we found long-range S1BF-spike to CA1-LFP synchronization to be homogeneous across inhibitory and excitatory cell types. These findings suggest distinct, cell-type contributions of low and high-frequency synchronization to intra- and inter-areal neuronal interactions.

## 1. Introduction

Cortical computation relies on the precise and flexible coordination of neuronal activity on multiple spatial and temporal scales. Rhythmic neuronal synchronization is a candidate mechanism for this coordination, creating an internal temporal reference frame that allows for the ordered activation of neurons at a specific time scale. Theoretical work indicates that slow and fast rhythms might subserve different functions in organizing neuronal communication in cortex, with slow rhythms synchronizing activity among distal neuronal groups, and fast rhythms synchronizing activity locally (Kopell et al., 2000). Here, we investigate how neocortical cells are entrained by oscillations at various frequencies within the local circuit and in connected areas, and how this depends on behavioral state and cell type. We address this question in rodent area S1BF, one of the main neocortical model systems for studying sensory processing, micro-circuit organization, and cortical plasticity (Brecht, 2007; Diamond et al., 2008; Fox, 2002; Petersen, 2007).

Transitions to active behavioral states are characterized by a loss of cortical synchronization at delta frequencies (<4 Hz) and associated with cortical theta, beta and gamma oscillations (Buzsáki, 2006; Harris & Thiele, 2011; McCormick et al., 2015). These oscillations can have a strictly local character, or can be coherent across brain areas, which may result from unidirectional entrainment, phase coupling between oscillators, or pacemaker activity (Akam & Kullmann, 2014; Buzsáki & Draguhn, 2004; Gielen et al., 2010; Kopell et al., 2000; Roberts et al., 2013; Steriade et al., 1993; Wang, 2010; Womelsdorf et al., 2014). There are ample examples of long-range beta (12–30 Hz) coherence between brain regions, which has been hypothesized to be a mechanism for top-down modulations (Bastos et al., 2015; Bressler et al., 2006; Brovelli et al., 2004; Buschman & Miller, 2007; Salazar et al., 2012). Theta oscillations (4–14 Hz) are prominent in hippocampus and prefrontal cortex, and implicated in the coordination of hippocampal with neocortical activity, subserving memory, decision making or navigation functions (Sirota et al., 2008; Womelsdorf et al., 2010). In sensory cortex, gamma-synchronization (30–90 Hz) is associated with local coding operations (Havenith et al., 2011; Vinck et al., 2010a; Womelsdorf et al., 2012), while inter-areal gamma coherence is thought to be involved in selective, feedforward routing (Bosman et al., 2012, 2014; Buschman & Miller, 2007; Gregoriou et al., 2009; Womelsdorf et al., 2007).

Few studies have performed simultaneous single unit and field recordings from multiple areas (i.e., spikes and fields in both areas) (e.g. Jia et al. (2013); Schomburg et al. (2014)), which is critical to interpret patterns of inter-areal coherence unambiguously (Buzsáki & Schomburg, 2015). This approach also allows one to compare the contributions of distinct neocortical cell types to local and inter-areal synchronization. It is important to understand these cell-type specific contributions because they determine the way in which synchronization governs neuronal interactions, and because they are informative about the mechanisms underlying rhythmic synchronization (Buzsaki & Wang, 2012; Cardin et al., 2009; Fries et al., 2007; Klausberger et al., 2003; Sohal et al., 2009; Wang, 2010; Womelsdorf et al., 2014). Here, we focus on LFPs and spike trains from different cell types recorded from S1BF, and relate these to simultaneous spike and LFP recordings from three other, connected brain areas (V1M, Perirhinal, CA1) in the awake, behaving rat. These have mono- and disynaptic connections with S1BF and contain neurons with tactile responses (Aronoff et al., 2010; Itskov et al., 2011; Iurilli et al., 2012; Naber et al., 2000; Paperna & Malach, 1991; Pereira et al., 2007; Vasconcelos et al., 2011). We study both intra- and inter-areal field-field and unit-field synchronization patterns, and examine how these depend on excitatory and inhibitory cell types and behavioral state. Using a whole-cell recording dataset, we examine the effects of state on gamma rhythmicity in membrane potential (*V_m_*) and validate our neuron type classification.

## 2. Material and Methods

We describe methods for two datasets separately: a dataset consisting of extracellular recordings obtained in rats and a dataset consisting of intracellular (whole-cell) recordings in mice. We refer to these two datasets as the *extracellular dataset* and the *intracellular dataset* throughout our manuscript.

### Extracellular dataset: Subjects

Data was collected from three 28–46 wk old male Lister Hooded rats (obtained from Harlan, Netherlands). During handling and behavioral training, animals were communally housed in standard cages under a reversed day/night cycle (lights off: 8:00 AM, lights on: 8:00 PM). During behavioral training and the main experiment, animals were food restricted to maintain their body weight at 85% of free-fed animals, taking the ad lib growth curves of Harlan and Rolls & Rowe (1979) as a reference (weights during recording were between 384–427 grams). From two days before surgery until after a full post-surgery recovery week, food was available *ad libitum*. Rats had *ad libitum* access to water during all phases of the experiment. After surgery, animals were housed individually in transparent cages (40 x 40 x 40 cm). All experiments were conducted according to the National Guidelines on Animal Experiments and were approved by the Animal Experimentation Committee of the University of Amsterdam.

### Extracellular dataset: Apparatus and stimuli

The animals were trained on a two-choice visual discrimination task set on a figure-eight maze (Figure 1) (114 cm x 110 cm). The paths of the maze were 7 cm wide (i.e., 7 cm separation between walls) and were flanked by walls that were 4 cm in height. The floor of the maze was elevated 40 cm above ground level. During the inter-trial-interval, the rat was confined to the middle arm of the figure-eight maze using two movable plexiglass barriers (Figure 1). Stimuli used were two equiluminant Wingdings (Microsoft, Redmond, WA) figures (either a diamond or plane; Figure 1) that had the same proportions of black and white pixels. The stimuli were simultaneously presented on two LCD (switch) monitors (Dell, 15 inch; Figure 1). At the sides of the maze arms into which the rat entered after the decision point, strips of rough (P40) or smooth sandpaper (P180) were attached to the inner walls of the maze (Figure 1). Reward pellets (BioServe, dustless precision pellets, 14 mg) were given in three ceramic white cups, with one cup in each arm (left and right) at the designated reward site, and one cup located in the “ITI confinement space”, close to the screens (referred to as the ‘front barrier’). After a correct trial, one pellet was given in the latter cup, at the beginning of each inter-trial interval. Eight photobeams were attached to the outer walls of the maze, with two photobeams in the middle arm, and three photobeams per side arm. The behavioral program was controlled using Matlab (Mathworks). Events that were detected by the behavioral apparatus or the commands issued by the behavioral program were directly time-stamped and synchronized with electrophysiological data by feeding them as inputs into the Neuralynx system.

**Figure 1:**
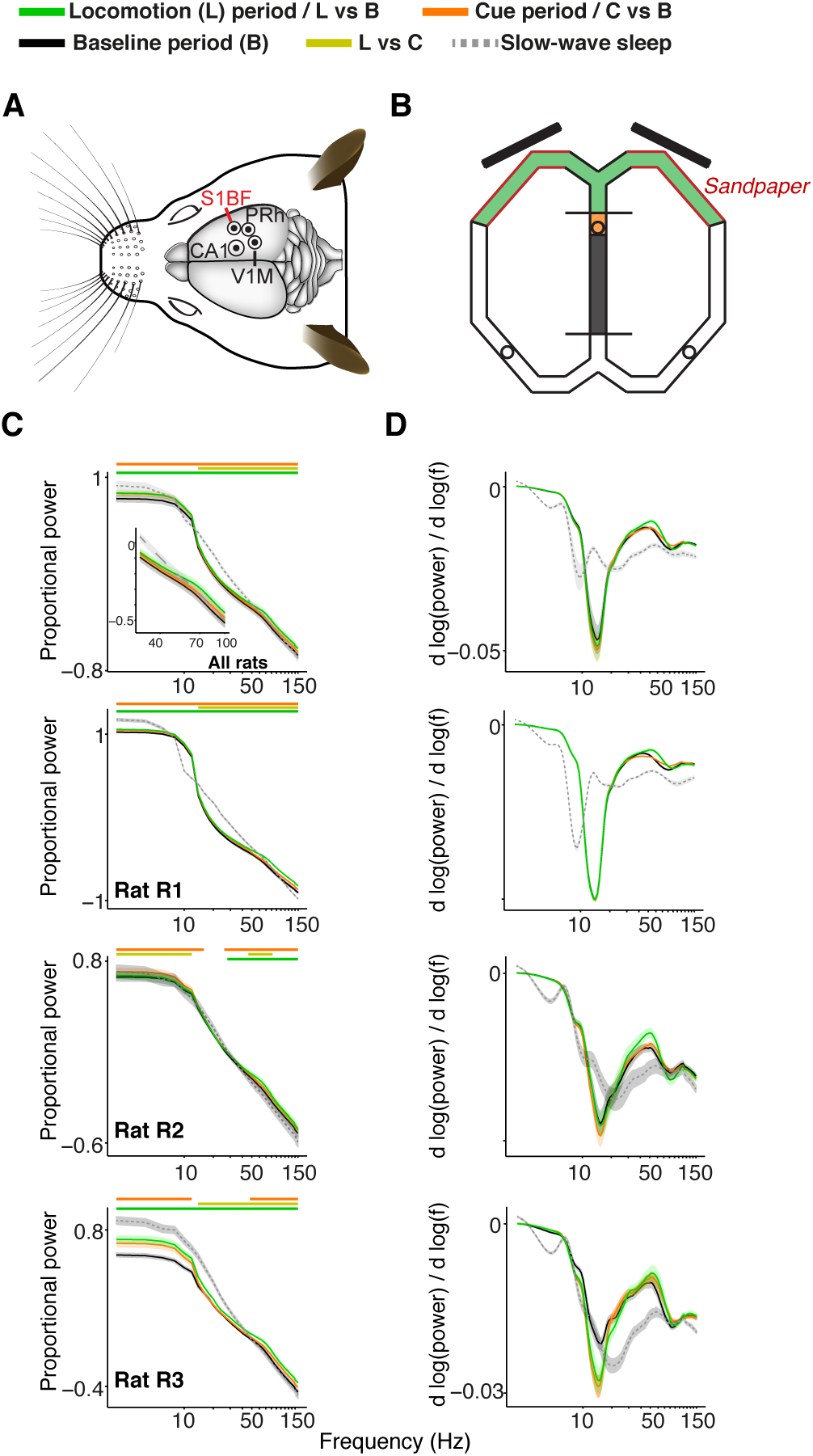
Recordings, task, and characteristics of power spectra for S1BF electrodes. **(A)** Overview of recordings. Craniotomies of the four brain areas are shown by circles, made for the target areas S1BF, dorsal CA1, perirhinal cortex and V1M. Eight recording (and one reference) tetrodes entered the brain per craniotomy. Adapted from (Paxinos & Watson, 2006). **(B)** Overview of the figure-8 maze. During the baseline period (grey), the rat is confined to the middle arm in the ITI confinement space by the two plexiglas barriers. Towards the end of the ITI, a sound indicates to the rat that the visual stimuli will appear if he breaks the infra-red photo-beam in the orange-shaded area. One second after breaking the infra-red beam, the S+ and the S-cue appear on the two 15 inch screens (black, oblique bars; onset of cue period, marked in orange). 4.2 s after stimulus onset the front barrier is removed (end of cue period) and the rat is allowed to make a response by entering either the left or the right arm. We refer to the 4.5–6.5 s period after cue onset as the locomotion period, when the rat approximately traversed the maze segments shown in green. The red walls close by the screens are covered by sandpaper cues which were indicative of the size of an upcoming reward. The circles show the locations of the three ceramic cups where the rat obtained his pellet rewards upon correct choices. **(C)** Average power spectra (normalized as explained in main text) as a function of frequency. Power spectra are shown on logarithmic (base 10) scale, and were, for all periods, normalized by dividing by the sum of power across frequencies for the baseline period. Inset shows data for frequency range of 30–100 Hz. Green, chartreuse and orange horizontal bars indicate significance of locomotion period relative to baseline, locomotion period relative to cue period, and cue period relative to baseline, respectively (p<0.05, Paired Rank-Wilcoxon Test, FDR correction for number of frequencies). **(D)** 1/*f* corrected power spectrum. This reveals that during wake states, (log) power decreased less steeply as a function of log_10_(*f*) around gamma frequencies, as compared to slow wave sleep states, which can also be seen from **(C)**. Shadings indicate SEM across sessions.

### Extracellular dataset: Training procedure

The two-choice visual discrimination task began with an inter-trial interval with random duration of 15–25 s, was followed by the onset of a 2 kHz sinusoid sound cue that lasted for 0.1 s, indicating that breaking the infrared photo-beam in the ITI space, close to the front barrier, would cause the visual stimuli to appear. Throughout all sessions performed by each rat, one of the two stimuli was designated the S^+^ stimulus, while the other stimulus was designated the S^−^. In every trial, both the S^+^ and S^−^ were presented, with the spatial location (right or left screen) of the stimuli varying randomly. 4.2 s after stimulus onset, the front barrier was removed manually. The rat could now enter one of the two side arms. The rat’s final choice was indicated by the rat breaking one of the two infrared photo-beams that were positioned beyond the visual screens at the end of the sandpaper walls (Figure 1; ‘Point of no return’). Hereupon, the stimuli were immediately turned off. We refer to these trials with late stimulus offset as ‘normal’ trials. We also included a set of ‘early-offset’ trials. During early-offset trials, stimuli were already turned off after 2 s. Upon correct choices, two or three pellets were manually placed in the ceramic cup that was located in the chosen arm. The conjunction of correct choice and rough sandpaper resulted in three pellets, while a correct choice and smooth sandpaper resulted in two pellets. When the rat entered the ITI space again, after making a correct choice (but not after an incorrect choice), an additional pellet was placed in the middle arm cup, and the front and back barriers were placed back in position, after which the next inter-trial-interval commenced. If rats attempted to walk back after passing the point of no return beam, the front barrier was placed on the maze, between the point of no return and reward cup, to block the rat’s return. Rough and smooth sandpaper were switched between arms every 10 trials. During recording sessions, rats performed 60.1 ± 18.7 (mean ± std) trials. It should be stressed that the questions addressed here do not focus on a detailed analysis of gamma-synchronization in relation to specific task components, such as the auditory, visual or tactile cues, nor on their integration. The analysis focuses instead on the features of gamma activity as it arises during naturalistic behavior in general, with special attention for task phases in which rats were locomoting versus largely immobile.

For the electrophysiological analysis, we only included trials in which the rat reached the reward port within 15s. Animals (rat R1, R2 and R3) performed respectively 70.21 ± 3.88 (mean ± SEM), 41.57 ± 2.1 and 44.54 ± 5.57 trials, at performances of 67.03 ± 1.24, 61.41 ± 2.75 and 60.59 ± 2.35%. We focused analysis in this paper on three different periods: locomotion (4.5 to 6.5 s after cue onset), baseline (-10 to 1 s before cue onset) and the cue period (0 to 4.2 s after cue onset). These periods had average locomotion velocities (across sessions) of respectively 25.73 ± 1.02, 2.44 ± 0.11 and 1.89 ± 0.09 cm/s.

### Extracellular dataset: Surgical procedure and recording drive

Each rat’s right hemisphere was implanted with a custom-built microdrive (Technology Center, University of Amsterdam) containing 36 individually movable tetrodes, including eight recording tetrodes directed to the visual cortex (V1M, -6.0 mm posterior and -3.2 mm lateral to bregma), eight to the dorsal hippocampal CA1 area (-3.5 mm posterior and -2.4 mm lateral), eight (and one reference) to the S1 Barrel Field (-3.1 mm posterior and -5.1 mm lateral) and eight to the perirhinal cortex (area 35/36, -5 mm posterior and -5 mm lateral), with one additional tetrode per area that could serve as a reference. Data were referenced against an electrode in the corpus callosum, unless stated otherwise. The microdrive’s design was based on a previously used split micro-drive (Lansink et al., 2007), weighed 23 grams and was 55 mm in height. The perirhinal bundle was placed at an angle of 17° with respect to a perpendicular orientation relative to the skull, such that tetrodes were aimed at area 35/36 (area border: -6 mm posterior, -6.4 mm lateral, and -6.2 mm ventral to bregma; Paxinos & Watson (2006)).

Prior to surgery (20–30 minutes), the rat received a subcutaneous injection of Buprenorphin (Buprecare, 0.01–0.05 mg/kg), Meloxicam (Metacam, 2 mg/kg), and Baytril (5 mg/kg). Rats were anesthesized using 3.0% (induction) and 1.0%-3.0% (maintenance) isoflurane. The rat was mounted in a stereotaxic frame. Body temperature was maintained between 35 and 36° C using a heating pad. After the cranium was exposed, six holes were drilled to accommodate surgical screws. Four holes (approximately 1.8 mm in diameter) were drilled for the four bundles holding the tetrodes. After removing the dura, the bundles were lowered onto the exposed cortex. We then fixed the whole microdrive to the skull and the surgical screws using dental cement. A skull screw located on the caudal part of the parietal skull bone contralateral to the drive location served as ground. After anchoring the drive, the tetrodes were lowered 0.4–1.0 mm (depending on the target area) into the cortex. Over the next seven days, the animal was allowed to recover, with *ad libitum* food and water available. The recording tetrodes were gradually lowered to their target region over the course of the first 7–9 days after implantation (using Paxinos & Watson (2006)), with electrode depths recorded daily. Depths were estimated by the number of turns of the guide screws, but also by online monitoring of the LFP and spike signals.

### Extracellular dataset: Histology

After the final recording session, current (12 *μ*A for 10 s) was passed through one lead per tetrode to mark the endpoint of the tetrode with a small lesion. The animals were deeply anesthetized with Nembutal (sodium pentobarbital, 60 mg/ml, 1.0 ml i.p.; Ceva Sante Animale, Maassluis, the Netherlands) and transcardially perfused with a 0.9% NaCl solution, followed by a 4% paraformaldehyde solution (pH 7.4 phosphate buffered). Following immersion fixation, transversal sections of 40 *μ*m were cut using a vibratome and stained with Cresyl Violet to reconstruct tetrode tracks and localize their endpoints. S1BF electrode positions were, on a daily basis, using both turning coordinates and electrophysiological signals, classified as superficial (L2–4), deep (L5–6), or ‘transition of L4–5’. An error margin of 250 *μ*m was used when assigning electrode positions to deep or superficial layers. Because gamma-synchronization was found in all layers, and our assignment of layers was rather coarse, we decided to pool data from different layers together.

### Extracellular dataset: Data acquisition and spike sorting

Using tetrodes (Gray et al., 1995) (nichrome, California Fine Wire, 16 *μ* per lead, gold-plated to 500–800 kΩ impedance at 1 kHz), we recorded neural activity with a 128-channel Digital Neuralynx Cheetah setup (Neuralynx, Bozeman MT). Signals were passed through a unity-gain pre-amplifier headstage, a 128-channel, automated commutator (Neuralynx, Bozeman, MT) and bandpass filtered between 600–6000 Hz for spike recordings. One ms epochs of activity from all four leads were digitized at 32 kHz if a signal on any of the leads of a tetrode crossed a pre-set voltage threshold. Local field potentials recorded on all tetrodes were continuously sampled at 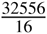 Hz, and bandpass filtered between 1–500 Hz. Spike trains were sorted to isolate single units using a semi-automated clustering algorithm followed by manual refinement (KlustaKwik, Ken Harris and MClust 3.5, A.D. Redish). During recordings, the rat’s behavior was videotracked at 25 Hz, and an array of light-emitting diodes on the headstage allowed offline tracking of the rat’s position. Automated and manual clustering of spikes was performed using the waveform peak amplitude, energy, and first derivative of the energy (energyD1 in MClust). Clusters were accepted as single units when having no more than 0.1 % of inter-spike intervals shorter than 2 ms.

### Extracellular dataset: Sleep epochs

Recordings of the task period were flanked by rest/sleep recordings of about 30–60 minutes. During these periods the rat was placed on a towel, folded into a flowerpot, which was located on a table placed above the maze. We identified slow wave sleep episodes by detecting the absence of movement and a high sharp wave ripple frequency in the hippocampus.

### Extracellular dataset: WPLI and power spectra

Data was analyzed in the ±10 s around stimulus onset, using custom-made Matlab and C software and the Matlab FieldTrip toolbox (Oostenveld et al., 2011). LFP-LFP coherence was computed using the WPLI (Vinck et al., 2011). For LFP-LFP coherence, we divided the data in segments of 0.5 s, and Fourier Transformed the LFP signals using multi-tapering tapering of 0.5 s windows, with spectral resolution of ±8 Hz. Denote the estimated cross-spectral density between two channels for the i-th segment (of length 0.5 s) out of *K* segments by *S* _12,_*_k_* (the ‘cross-spectrum’). The WPLI (weighted phase lag index) (Vinck et al., 2011) is a measure of phase-synchronization that is not spuriously increased by volume conduction and has been argued to have reduced noise sensitivity relative to previous measures of phase-synchronization that utilize the imaginary part of the cross-spectrum (Nolte et al., 2004; Stam et al., 2007). In addition, we showed that even in case of two dependent (interacting) sources and sensors, the position of the sources relative to sensors (i.e., the specific volume conduction mixing coefficients) does not alter the WPLI (Ewald et al., 2012; Vinck et al., 2011). The debiased WPLI estimator is defined as and is a nearly unbiased estimator of the square of the WPLI statistic, which is defined as 

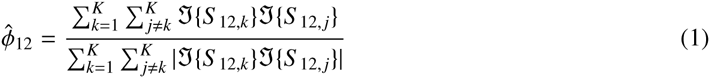
and is a nearly unbiased estimator of the square of the WPLI statistic, which is defined 

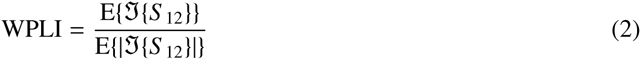
where *S*_12_ is a random variable identically distributed to *S* _12,_*_j_* for all *j*, E{} is the expected value operator, and ℑ denotes imaginary component. Note that a direct estimate of the WPLI is strongly biased by sample size, and that the debiased WPLI is a nearly unbiased estimator of the squared WPLI. The (debiased) WPLI estimates were then averaged across all channel-combinations.

To visualize what band-limited LFP-LFP phase-synchronization as measured in the frequency domain corresponds to in the time domain, we computed an inverse Discrete Fourier Transform (DFT) of the spectral coherency function, and its imaginary component, which is not spuriously increased by volume conduction of a single source (Nolte et al., 2004). The spectral coherency was estimated as 

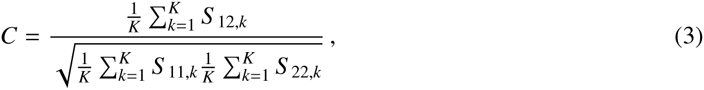
where *S*_11,_*_k_* and *S*_22_,*_k_* are the estimated power spectra for the 1st and the 2nd signal for the kth segment. We then computed the inverse DFT of the coherency function, and band-pass filtered it between 30 and 120 Hz. This gives similar information as the cross-correlation function, except that the cross-correlation function emphasizes the contribution of spectral components with high power, whereas the inverse DFT of the coherency function emphasizes the contribution of spectral components with high predictive power (of one signal by the other signal) (Wiener, 1956).

For the power spectrum, we also used the same settings for spectral estimation as for the coherence analysis. Similar to (Vinck et al., 2013), we normalized the raw power spectra by dividing the raw power at each frequency by the mean power across frequencies for the baseline period, and taking the log_10_ transform (Figure 1), i.e.

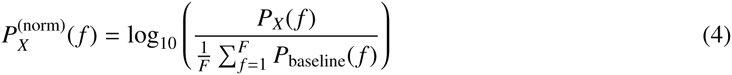
where *P_X_*(*f*) is the raw power in a period *X*, *P*_baseline_ the raw baseline power, *f* is frequency and *F* the number of frequencies. Thus, for all behavioral periods (cue, locomotion, and baseline), the power spectrum was normalized to the total power in the baseline condition. This normalization procedure does not change the shape of the power spectrum, and has two main advantages: a) Because we normalize the power spectrum in all behavioral conditions relative to one single condition (baseline), we can compare differences in raw power between conditions. b) Because the power is normalized per session before averaging, it squeezes out the variance driven by global changes in signal energy across sessions and animals.

We also computed the slope of the normalized power spectrum as

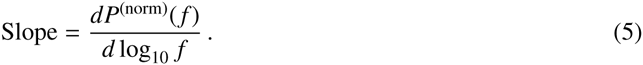

Note that computing this slope amounts to performing a correction for the linear trend in log(power) vs log(frequency), since if log(power) is a linear function of log(frequency), the slope is a constant negative.

### Extracellular dataset: The spike-LFP pairwise phase consistency (PPC)

For each frequency *f*, we determined the spike-LFP phases by cutting out LFP segments of length 5/*f* s (i.e., 5 cycles) centered around each spike. Spikes were only related to LFPs recorded from a different electrode to avoid contamination of the LFP by the spike itself. For spikes that fell around the border of an analysis window, we determined the phase of the LFP by cutting out an LFP segment that started at the border of the window, i.e. not centered around the spike. The spike-LFP phases were then obtained as the complex arguments of the Kaiser (with *β* = 9) tapered LFP segments. With a Kaiser window, a 50 Hz LFP signal results in -10 dB energy (from leakage) at 30 and 70 Hz. We always averaged the spike-LFP phases across the different electrodes (excluding the electrode on which the unit under consideration was recorded) before computing measures of phase consistency.

The strength of spike-LFP phase locking was quantified by the PPC, which is unbiased by the number of spikes (Vinck et al., 2012) (see also van Wingerden et al. (2012); Vinck et al. (2013)). For the *j*-th (or *l*-th) spike in the *m*-th (or *k*-th) trial we denote the spike-LFP phase as *θ_m,j_* (or *θ_l,k_*), where dependence on frequency is omitted in what follows. The PPC is then defined as

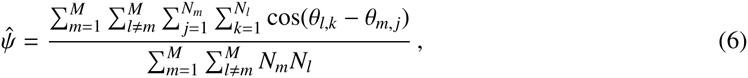
where *N_m_* is the number of spikes for the *m*-th trial. The PPC quantifies the average similarity (i.e., in-phaseness) of any pair of two spikes from the same cell in the LFP phase domain. Note that all pairs of spikes from the same trial are removed by virtue of letting *l ≠ m* in eq. 6, because spike phases from the same trial can typically not be treated as statistically independent random variables (Vinck et al., 2012).

To test whether differences in PPC values across frequencies were significant across conditions, we used multiple comparison corrected randomization testing according to Korn et al. (2004). This randomization testing works in a sequential fashion: We compare whether the largest observed difference in PPC values across conditions exceeds the 97.5% percentile of the largest observed differences in PPC values in the randomization distribution. This randomization distribution was generated by randomly shuffling the condition labels across cells or LFPs, without replacement. In case of testing differences between cell types, we randomly shuffled the cell type labels across cells (without replacement). Significant frequencies are then identified in a step-wise fashion. The advantage of this procedure relative to cluster-mass based randomization testing, which is popular in electrophysiology (Maris et al., 2007), is that we do not put low frequencies to a disadvantage because they have smaller bandwidths. Furthermore, our procedure allows one to detect multiple significant clusters, while the cluster-mass based procedure of Maris et al. (2007) does not.

To test whether inter-areal PPC values were significantly greater than zero, we again used randomization testing. For each permutation, we shuffled spikes and LFPs and recomputed PPC values. We then used the randomization testing procedure of Korn et al. (2004) to see at which frequencies the average PPC exceeded the PPC of the randomization distribution.

### Intracellular dataset: Intracellular whole-cell recordings in vivo

Full methods of analyzed data have been reported in (Gentet et al., 2010, 2012). In brief, whole-cell patch clamp recordings were performed in the C1 column of barrel cortex in 80 awake, head-restrained transgenic male mice (6–10 weeks; GAD67, GIN and Sst-Cre lines) as previously described (Gentet et al., 2010, 2012). Analyzed traces are issued from previously published datasets (Gentet, 2012; Gentet et al., 2010) as well as yet unpublished datasets. In brief, intracellular membrane potential dynamics during sequences of free whisking in air and active touch sequences (average rate of touches = 6.3 Hz) of at least 2 s durations were compared with closely time-spaced sequences of quiet wakefulness for each cell to obtain cell-type specific correlates of gamma-band oscillations. We used the following procedure to compare *V_m_* fluctuations between conditions:

1) We removed the action potentials ([-0.8 to 3 ms]) from the *V_m_* traces. 2.) We always compared segments of 500 ms of whisking and baseline episodes with the same number of spikes. Despite stratifying the number of spikes per segment, significant *V_m_* differences between whisking and baseline episodes were retained, indicating that our stratification procedure did not remove all meaningful dynamics from the voltage membrane potential. 3.) We then computed the Lomb-Scargle periodogram which amounts to estimation of a power spectrum for time series with missing values using least-squares fits of sine and cosine functions. This method, commonly used in astrophysics, is based on the same basic idea as Fourier analysis but does not require regular sampling, and it does not give rise to artifacts that can arise from interpolating the missing values (Ruf, 1999). Thus, we computed the spike count per 500 ms segment, and for each unique spike count that was observed in both conditions (e.g., segments of zero spikes in both conditions), we estimated the power spectrum using the Lombe-Scargle periodogram by averaging across all segments (i.e., all segments of zero spike counts). These estimates were then averaged across all unique spike counts occurring in both conditions (e.g., segments of zero, one and two spikes). 4.) As a first control, we used Brownian noise 1/*f*^2^ data to verify that our procedure did not lead to any biases. 5.) For the pyramidal cells, which have low firing rates but longer action potential durations, we also removed -2 to 10 ms around each spike as a control.

## 3. Results

The Results section is organized as follows. We first focus on local synchronization within area S1BF, subsequently analyzing univariate LFP signals, coherence between S1BF LFP signals, and phase locking between spikes fired by different cell types and LFPs. We then proceed with an analysis of inter-areal synchronization patterns. We finish with an analysis of the intracellular dataset, containing whole-cell recordings from specified excitatory and inhibitory cell types in area S1BF of awake, head-fixed mice. This latter dataset extends and complements the analysis of spike-field phase locking using the extracellular dataset.

### Extracellular dataset: LFP power analysis and LFP-LFP phase-synchronization

We recorded spikes and LFPs from 32 tetrodes in three awake behaving rats performing a left-right discrimination task on a figure-8 maze (Figure 1A-B; N=18, 15, and 13 sessions for rats R1, R2 and R3, respectively). Simultaneous recordings were made from four separate tetrode bundles, each containing eight tetrodes that were horizontally separated by 100–1000 *μ*m. Tetrodes from the four bundles were positioned in different brain areas, namely S1BF, the dorsal CA1 field of the hippocampus, perirhinal cortex (area 35/36), and the monocular field of the primary visual cortex (V1M). Data within each of the areas was always rereferenced to the local average, unless explicitly mentioned otherwise, as this suppresses (but does not fully eliminate) the contribution from distal current sources.

S1BF LFP power spectra exhibited two features: (1) power roughly fell off at a rate of *1*/*f^α^* (Figure 1C), which is generally characteristic for EEG signals (Buzsáki & Draguhn, 2004), and (2) a peak around gamma frequencies in the 1/*f* corrected power spectrum, which corresponds to a convexity in the power spectrum (Figure 1D). Gamma power and gamma power slope were significantly increased during the locomotion period (4.5 to 6.5 s after cue onset) relative to the baseline (-10 to 1 s before cue onset) and the cue period (0 to 4.2 s after cue onset; Figure 1C; p<0.05, FDR correction for multiple comparisons, Rank-Wilcoxon Test). This increase in power was accompanied by an increase in peak gamma frequency. The gamma peak in the 1/*f* corrected power spectrum was strongly quenched during slow wave sleep episodes, in which the log power spectrum was a near linear function of log frequency around gamma frequencies (Figure 1C-D). Because power, LFP-LFP and spike-LFP phase-synchronization spectra were highly similar in the cue and the baseline period, we restrict the comparison of states mainly to the baseline and locomotion period in what follows.

### Extracellular dataset: Phase synchronization between S1BF LFPs

Next, we asked to what extent neuronal activity, as measured through population mass LFP signals, was coherent across sites separated by 100–1000 *μ*m within area S1BF. We observed prominent band-limited gamma phase-synchronization between barrel LFPs, with spectral peaks around 60-70 Hz (Figure 2A-C). Phase-synchronization was measured using the WPLI (Weighted Phase Lag Index), which benefits from reduced noise sensitivity and is not spuriously increased by (instantaneous) volume conduction of single current sources or the use of a common reference (WPLI; Vinck et al. (2011)). Thus, volume conduction of single current sources to multiple S1BF channels is unlikely to explain the observation of band-limited gamma phase-synchronization between S1BF electrodes. S1BF gamma phase-synchronization was also observed when it was indexed by a standard LFP-LFP phase locking statistic (Pairwise Phase Consistency, Vinck et al. (2010b)) which, as opposed to WPLI, does take the real part of the cross-spectral density (i.e., instantaneous interactions) into account, although it showed a uniform increase in phase-synchronization across frequencies (Figure 2B).

**Figure 2:**
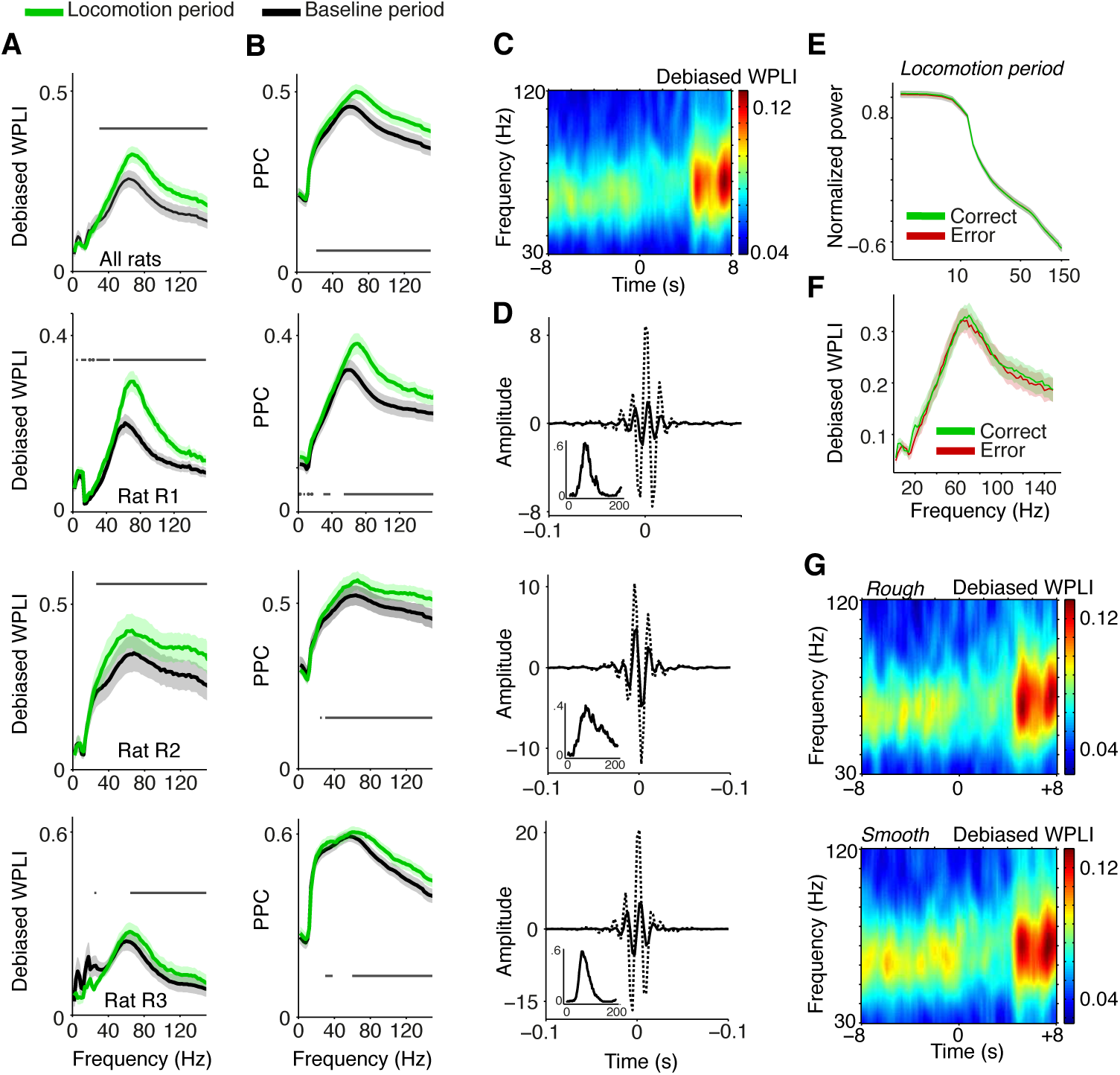
Phase synchronization between S1BF LFPs. **(A)** Average LFP-LFP phase synchronization [debiased WPLI, Vinck et al. (2011)] between S1BF LFPs, separately for the baseline (-10 to 1 s relative to cue onset), and locomotion (4.5 to 6.5 s after cue onset) period. S1BF electrodes had a horizontal separation of about 100 - 1000yum. Shadings indicate SEMs across sessions. Horizontal bars indicate significance of locomotion period relative to baseline (p<0.05, Rank-Wilcoxon Test, FDR correction for number of frequencies). Second to fourth row show individual rats R1 (N=18 sessions), R2 (N=15 sessions) and R3 (N=13 sessions). **(B)** Same as **(A)**, but now quantifying LFP-LFP phase synchronization with the PPC (Pairwise Phase Consistency). **(C)** Time-frequency representation of debiased WPLI around visual cue onset at t=0. For this analysis wavelets with 9 cycles per frequency and Hanning tapers were used. Gamma phase-synchronization is found in all behavioral periods, but increases during locomotion period. **(D)** Inverse Discrete Fourier transform (DFT) of coherency (dashed) and imaginary part of the coherence (solid) for example electrode pairs in rat R1, R2 and R3, filtered between 30 and 120 Hz. This depicts the time-domain representation of the spectral coherency and reveals oscillatory side-lobes, indicating that the observed gamma phase-synchronization indeed had a quasi-periodic nature. Inset shows debiased WPLI as a function of frequency, for this example pair. **(E)** Normalized power, as in Figure 1, for the locomotion period, and correct and incorrect trials. **(F)** Average LFP-LFP phase synchronization between S1BF LFPs, for the locomotion period, and during correct and incorrect trials. **(G)** As **(C)**, but now for trials in which the walls were covered with rough (top) and smooth (bottom) sandpaper.

To visualize what the frequency-domain LFP-LFP phase-synchronization spectra correspond to in the time domain, we computed the inverse Discrete Fourier Transform (DFT) of the spectral coherency and its imaginary component, which is not spuriously increased by volume conduction (Nolte et al., 2004) (see Material and Methods). The inverse DFT of the coherency function revealed oscillatory side-lobes for all three rats, showing that the observed gamma phase-synchronization indeed had a rhythmic nature (Figure 2D).

Gamma peaks in LFP-LFP phase-synchronization spectra were observed across all behavioral periods; thus, they occurred under a diverse range of naturalistic behavioral conditions. Gamma phase-synchronization was increased during the locomotion period in comparison to the baseline period (Figure 2A-C, p<0.05, FDR multiple comparison correction, Paired Rank-Wilcoxon Test), consistent with the power spectrum analysis.

We did not observe significant differences in gamma-synchronization or gamma power between correct and incorrect trials (Figure 2E-F) and trials in which the walls were covered with rough or smooth sandpaper (Figure 2G). In summary, the occurrence of gamma activity is supported by band-limited gamma phase synchronization between S1BF LFPs and shows a significant dependence on behavioral state.

### Extracellular dataset: Classification of cells according to waveform and firing characteristics

We now come to our main question, namely how distinct S1BF cell types are entrained by rhythmic fluctuations at various frequencies within the local area and in connected areas. Before analyzing the patterns of phase locking of 469 isolated single units from area S1BF (N=282, N=70 and N=117 cells in rats R1, R2 and R3, respectively), we first classified cells into GABAergic FS cells and multiple classes of excitatory cells. There exist three main classes of GABAergic cells in rodent S1BF (Gentet et al., 2012; Rudy et al., 2011): PV (Parvalbumin-positive), SSt (Somatostatin-positive) and NFS (Non-fast spiking) cells. PV and SSt cells account for approximately 70% of GABAergic cells across rodent S1 layers (Rudy et al., 2011). Intracellular recordings in L2/3 S1BF in awake mice have shown that both SSt and PV cells have narrow waveforms, whereas excitatory cells and NFS GABAergic cells have broader waveforms (Gentet et al. (2010, 2012), see also Cardin et al. (2009); McCormick et al. (1985) and our Figure 10 in which we reanalyze the data of Gentet et al. (2012)).

Also in our S1BF tetrode recordings, the distribution of peak-to-trough waveform durations was significantly bimodal (Figure 3A-B; p<0.05, Hartigan’s dip test). We divided the class with narrow waveforms into units with fast waveform repolarizations (FS), and units with slow waveform repolarizations (unclassified cells (UC); Figure 3A-B). FS cells (N=45) had higher firing rates than UCs (N=33) and cells with broad waveforms (N=391) (Figure 3C; p<0.001 for all comparisons among UCs, FS and cells with broad waveforms, Rank-Wilxocon Test), in agreement with the intracellular dataset (see below). We therefore assume that the FS class is the union of the PV and SSt cell classes, whereas the cells with broad waveforms correspond to the excitatory cell class, although a small percentage of NFS GABAergic cells may have been included in this group; we henceforth refer to the broad spiking cells as E cells, and note that these are only putatively identified as excitatory. FS and E cells were typically recorded from superficial/granular layers (FS cells: L5/6: 4; L4/5 transition: 10, L2–4: 31; E cells: L5/6: 34, L4/5 transition: 93, L2–4: 264; n.s. for all FS vs. E comparisons for all layers, randomization test).

**Figure 3:**
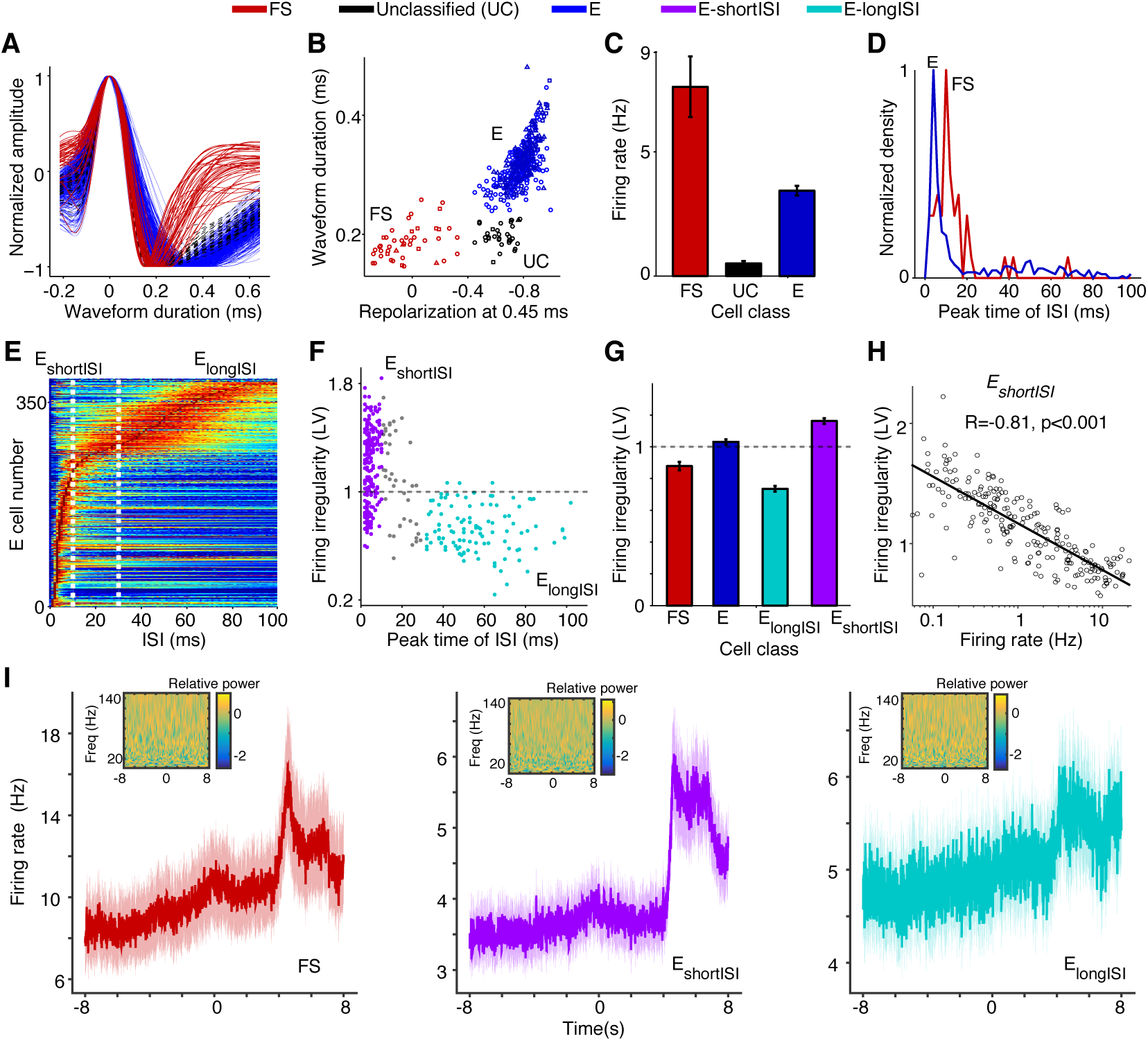
Assignment of (extracellularly recorded) S1BF units to electrophysiological cell classes. **(A)** Normalized waveform amplitude as a function of time (ms) from AP peak, separately for FS (fast spiking), E (putative excitatory) and UC (unclassified) cells. **(B)** Normalized AP amplitude at 0.45 ms (repolarization) vs. waveform peak-to-trough duration (ms), revealing three electrophysiological cell classes. Circles, squares and triangles correspond to Rat 1, 2 and 3, respectively. **(C)** Mean ± SEM of firing rate (Hz) for different cell classes. **(D)** Distribution of ISI (Inter Spike Interval) peak times across E and FS cells. **(E)** ISI histograms for all E neurons. The data in each row were normalized by dividing by the maximum (color scale ranges from 0 to 1). **(F)** Peak time of ISI vs. firing irregularity, which was measured using the LV (Local coefficient of Variation, Shinomoto et al. (2009)). E_shortISI_ cells were defined to have ISI peak times shorter than 10 ms. E_longISI_ cells were defined to have ISI peak times in excess of 30 ms. Gray dots correspond to cells attaining intermediate values. **(G)** Mean ± SEM of firing irregularity (LV) for different cell classes. **(H)** Log_10_ of firing rate vs. firing irregularity (LV), together with linear regression fit. Each dot corresponds to one E_shortISI_ cell. **(I)** Average peri-stimulus histograms ± SEM (bin size 10 ms) as a function of time from visual cue onset (in seconds), separately for FS, E_shortISI_ and E_longISI_ cells. Insets show the wavelet transform of the average peri-stimulus histogram with bin size 2 ms, using 7 cycle wavelets with Hanning taper for each frequency. Relative power (i.e. power normalized to total sum power), obtained via wavelet transform, is shown on log_10_ scale.

Based on anatomical data, we estimate the NFS GABAergic group to constitute only a minor <1–2% of all recorded cells in the E class, as they constitute only 30% of interneurons, out of which 50% reside in L1 (Beaulieu, 1993; Rudy et al., 2011), from which we did not make recordings. The observed differences between FS and E cells in terms of waveform characteristics and their observed percentages (9.6% FS cells and 90.4% putative excitatory cells) agree with data from L5 of S1Tr (Bartho et al., 2004), L2–4 of S1BF in mice (Cardin et al., 2009), and anatomical studies (Beaulieu, 1993; Lefort et al., 2009). UCs, which are presumably excitatory given their low firing rates, were not included in further analyses.

To identify excitatory cell classes, we computed two statistics on the cells’ spike trains. First, for each cell we computed the peak time in the ISI (inter-spike interval) histogram (Figure 3D-F). Second, for each cell we quantified the local coefficient of variation (LV), a measure of firing irregularity (Figure 3F-G). The LV measures the variability of subsequent inter spike intervals with limited influence of firing rate non-stationarities (LV=1, LV>1 and LV<1 indicate Poisson-like, irregular firing and regular firing, respectively). We found that, in contrast to FS cells, excitatory cells were a highly heterogeneous group. For E cells, the distribution of ISI peak times was bimodal (Figure 3D-F; Hartigan’s dip test, P<0.05). We could thus identify two classes of putative excitatory cells: cells with early ISI peaks having many short ISI intervals (named E_shortISI_) and cells that had late ISI peaks having many long ISI intervals (named E_longISI_; Figure 3D-F). These two extreme firing behaviors were previously identified by Bartho et al. (2004) in L5 of S1Tr (S1 Trunk region). E_longISI_ cells were more likely to be recorded from superficial layers (L5/6: 7, L4/5 transition: 12, L2–4: 95) than E_longISI_ cells (L5/6: 27, L4/5 transition: 71, L2–4: 138; p<0.05 for all comparisons between E_shortISI_ and E_longISI_ cells across all layers, randomization test). The E_shortISI_ cells showed substantial variability in firing irregularity, with a significantly bimodal distribution of LV values (p<0.05, Hartigan’s dip test). We found that this variability in firing regularity (i.e. LV) showed a very strong correlation to unit firing rate: E_shortISI_ cells with high firing rates tended to be non-bursty, whereas irregularly firing E_shortISI_ cells had low firing rates (Pearson’s r of log(firing rate) x LV = 0.81, p<0.001, Figure 3H). We refer to cells having a combination of irregular firing and an early ISI peak as ’bursting’.

We found that the firing rates of the three main cell classes tended to increase during the locomotion period in comparison to the baseline and the cue period (Figure 3I). Such a rate increase is consistent with previous reports on the dependence of firing rates on locomotion in both V1M and area S1 (McGinley et al., 2015; Niell & Stryker, 2010; Reimer et al., 2014; Vinck et al., 2015). We found that the increase in firing with locomotion was particularly pronounced for E_shortISI_ and FS cells, while it was less pronounced for E_longISI_ cells. We also computed the wavelet transforms of the average peri-stimulus histograms in order to examine whether the evoked firing responses contained energy in a particular frequency band. We found that the energy of the average peri-stimulus histogram was uniformly distributed over frequencies. Based on this finding, we do not expect rhythmicity of neuronal firing simply because of evoked activity.

In summary, the recordings in S1BF allowed us to distinguish multiple inhibitory and excitatory cell classes on the basis of extracellular waveform and firing rate statistics.

### Extracellular dataset: Cell-type specific spike-LFP phase locking

We then proceeded with the analysis of the phase locking of these cell types to local S1BF LFP fluctuations. The goal of this analysis was to gain further insight into the cellular basis of local gamma synchronization and its dependence on behavioral state. We related spikes from isolated single units to all LFPs recorded simultaneously from eight separate electrodes, excluding spike-LFP combinations from the same electrode. The precision of spike-LFP phase locking was quantified using the spike-LFP pairwise phase consistency (PPC), a metric unbiased by spike count and spike train history effects (Vinck et al., 2012). Figure 4 shows an example of an FS cell with a characteristically strong locking to gamma fluctuations in the LFP signal.

**Figure 4:**
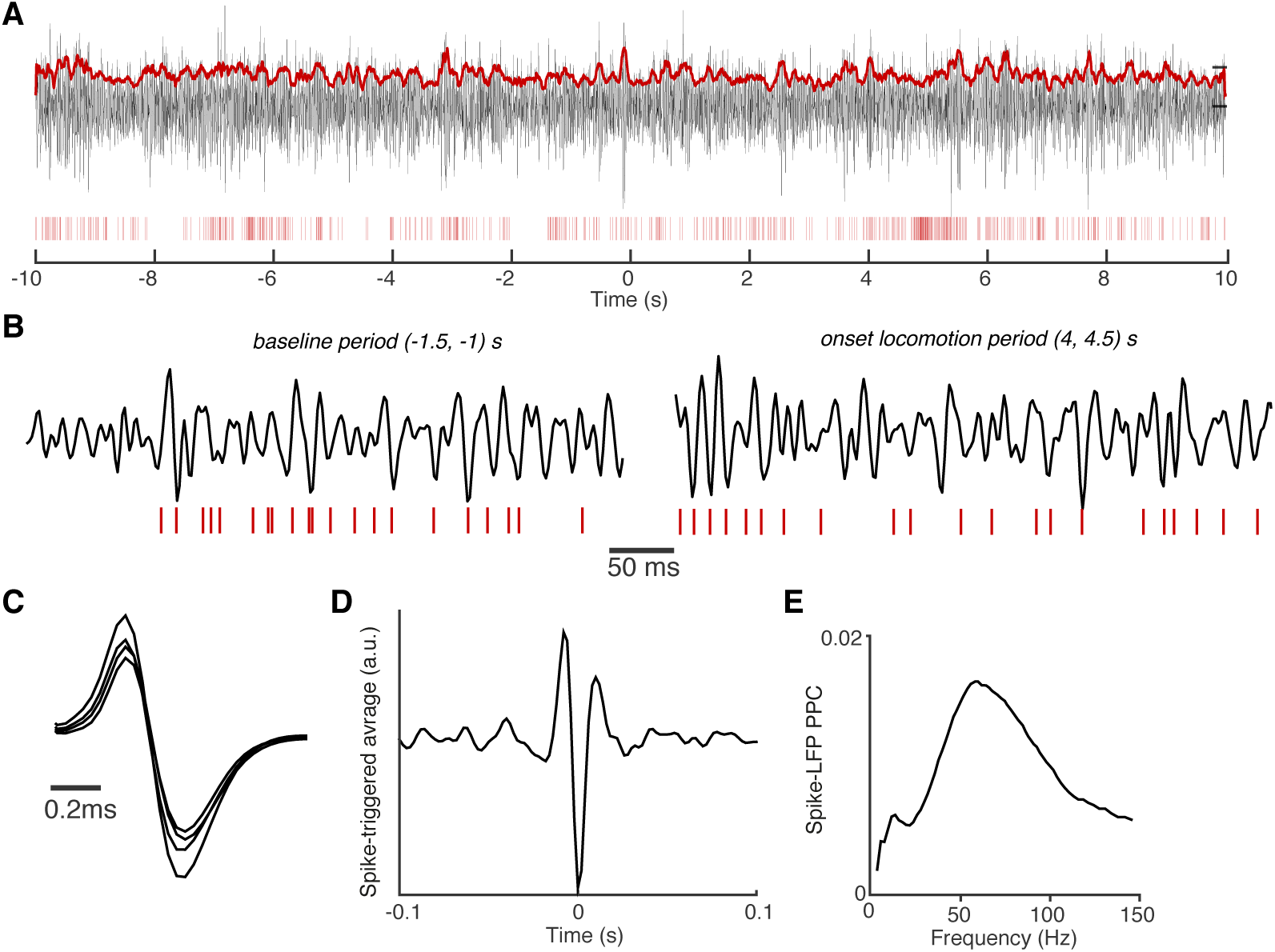
Example of spike-field coupling in area S1BF. **(A)** Example trial with spike trains of an S1BF FS cell (red vertical bars) and filtered S1BF LFP trace (30–120 Hz). LFP was recorded from another tetrode than the FS cell, positioned in the same layer and was roughly 0.4 mm away. Both electrodes were positioned in layers 2/3. LFP has trace plotted in arbitrary units. Red continuous trace: the 100 ms average of the Hilbert amplitude envelope of the filtered LFP signal. Note that the FS fires at high rates that are characteristic for GABAergic interneurons. t=0 corresponds to onset of visual cue. **(B)** Shown are different periods of trace in **(A)** with higher temporal resolution. All LFPs on the same scale (arbitrary units). **(C)** Average spike waveforms for the FS cell. Note the different waveform amplitudes recorded on the four different leads of the tetrode. **(D)** Average spike-triggered-average LFP over entire task period, filtered in 30–120 Hz band (arbitrary units). The spike-triggered-average LFP decorrelates quickly over time, reflecting the broad-band nature of spike-LFP locking. **(E)** Spike-LFP PPC spectra for the FS cell measured entire task period.

Average phase locking spectra of E and FS cells showed two visible peaks, one in the beta (12–20 Hz) and one in the gamma range (40–90 Hz). Beta locking was stronger in FS than in E cells, and stronger in E_shortISI_ than in E_longISI_ cells (Figure 5A-B). FS cells also exhibited prominent gamma peaks in PPC spectra and had higher gamma phase locking than E cells (Figure 5A, p<0.05, randomization test, multiple comparison correction [MCC] using Korn et al. (2004)). This finding was consistent across behavioral periods and when pooling behavioral periods together (Figure 5A, left panel). Gamma phase locking did not differ between behavioral states for FS cells, but was significantly higher for E cells during the locomotion period than in the baseline period (MCC randomization test). Thus, the increase in LFP-LFP gamma phase-synchronization and LFP power that was observed during the locomotion period co-occurred with an increase in E cell gamma locking, while FS gamma locking tended to be stable across behavioral periods. Both E_shortISI_ and E_longISI_ cells showed a gamma peak during the locomotion period, but gamma locking was stronger for EshortISI than E_longISI_ cells in this period (Figure 5B; MCC randomization test).

**Figure 5:**
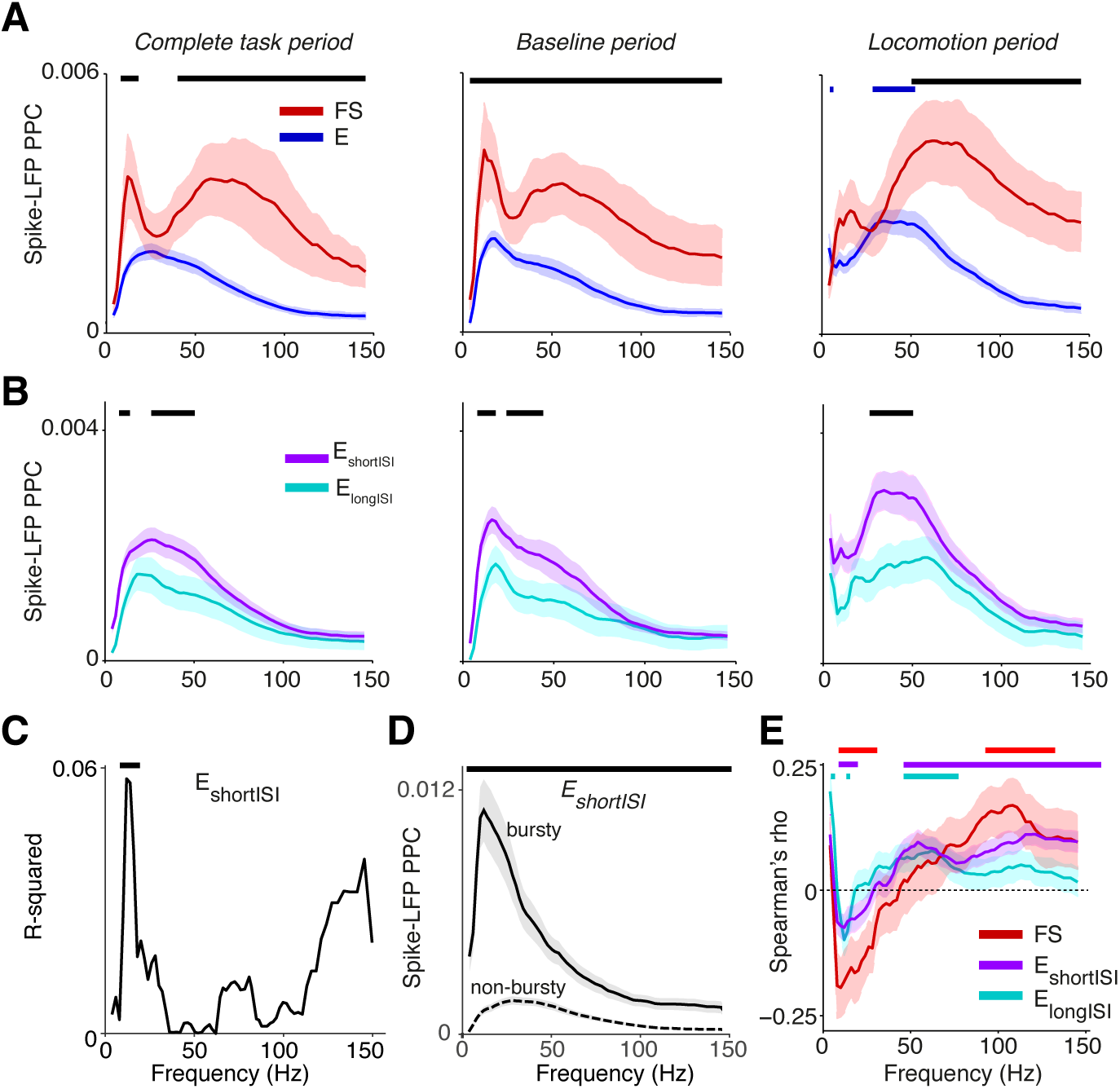
Phase locking of S1BF cells to S1BF LFPs. **(A)** Mean ± SEM of spike-LFP PPC (Pairwise Phase Consistency) as a function of frequency (Hz), separately for FS and E cells. Left: PPCs were computed over all recorded spikes in a window of ± 10 s around cue onset. Middle and right column correspond to baseline and locomotion period, respectively. Black significance bars indicate significant differences between FS and E cells. Blue bars indicate significance of phase locking difference relative to baseline period for E cells (multiple comparison corrected randomization test, Korn et al. (2004)). Besides a gamma peak, the PPC spectra show beta peaks around 12–20 Hz. **(B)** Same as **(A)**, but now for E_longISI_ and E_shortISI_ cells. **(C)** Increase in R-squared (explained variance) by adding the LV to a multiple regression model, in which we predict PPC values (measured in entire task period) from LV and log_10_ firing rates for E_shortISI_ cells. Black bars indicate significance of regression coefficient (p<0.05, FDR corrected for number of frequencies). **(D)** Mean ± SEM of PPC spectra (measured in entire task period) for bursty E_shortISI_ cells and non-bursty firing E_shortISI_ cells. We identified these cells using fuzzy c-means clustering on the distribution of LV values, with 0.95 probability membership cut-offs. Black bars indicate significance of phase locking difference (multiple comparison corrected, Korn et al. (2004)). **(E)** Spearman’s *ρ* between a cell’s average spike density at time *t*, computed in a window of 2 s, and its PPC value at time *t*, across time-points. This correlation was computed for each cell separately, taking the -10 to 10 s period relative to cue onset. Spearman’s *ρ* values were then averaged across cells. Shown are means ± SEMs. Horizontal bars indicate significance at p<0.05 for each cell class.

Within the E_shortISI_ group, we found that cells that were more irregularly firing (i.e., irregularly bursty with a high LV) tended to show stronger beta phase locking (Figure 5C). To illustrate the difference in beta phase locking between non-bursty and bursty E_shortISI_ cells, we clustered the E_shortISI_ cells in two groups, using fuzzy c-means clustering (cut-offs at 0.95 membership probability). This analysis revealed strong beta phase locking in bursting E_shortISI_ cells, whereas the phase locking spectrum of non-bursty E_shortISI_ cells was dominated by gamma-frequencies (Figure 5D). Together, these findings suggest that beta rhythmicity arises from an interaction between FS cells and irregularly bursty cells.

We investigated correlations between firing rate and phase locking by computing, for each cell, the mean spike density and mean PPC at every point in time during the ± 10 s around cue onset (using a window of ±1 s). We found that firing rates were positively correlated with gamma PPCs for both E_longISI_ and E_shortISI_ cells, while they were positively correlated with supra-gamma (100–120 Hz) PPCs for FS cells (Figure 5E). Beta phase locking (12–20 Hz) was negatively associated with firing rates of all cell types. Thus, gamma and beta oscillations tended to be associated with network activations and deactivations, respectively.

We conclude that there is rhythmic entrainment of inhibitory and excitatory neurons in barrel cortex both at slower (beta) and faster time scales (gamma). This phase locking is cell type specific, and it shows a dependence on behavioral state in case of excitatory, but not FS cells.

### Extracellular dataset: Cell-type specific spike-LFP phases

The way in which rhythmic synchronization shapes local interactions depends on the precise temporal patterning of distinct neuron types. The order in which different cellular phenotypes fire is also informative about the local mechanisms contributing to oscillations (e.g., ING and PING mechanisms; Bartos et al. (2007); Börgers & Kopell (2005); Buzsaki & Wang (2012); Csicsvari et al. (2003); Eeckman & Freeman (1990); Tiesinga & Sejnowski (2009); Wilson & Cowan (1972)). We therefore examined the temporal order in which distinct S1BF cell classes fired relative to S1BF oscillations. Here, we did not use a local reference but used a reference in the corpus callosum as this does not cause a phase shift in the local phase angle. Both FS and E cells fired on average around the trough of the LFP gamma cycle (for complete task period: at peak gamma frequency 66 Hz, FS: mean ± 95% c.i. = 175.0 ± 25.5°, E: 176.9 ± 11.0°, Figure 6A), in agreement with (Siegle et al., 2014). The confidence levels indicate that the delay between FS and E cells was restricted within roughly 1 ms, a delay that is too short to result from excitatory feedback. The absence of a delay between FS and E cells was also observed at other frequencies, and was consistent across several behavioral periods (Figure 6A, n.s. at all frequencies for all behavioral periods, Circular ANOVA). We did not detect a significant gamma phase difference between E_shortISI_ and E_longISI_ cells (Figure 6B, n.s. at frequencies between 40 and 100 Hz, Circular ANOVA).

**Figure 6:**
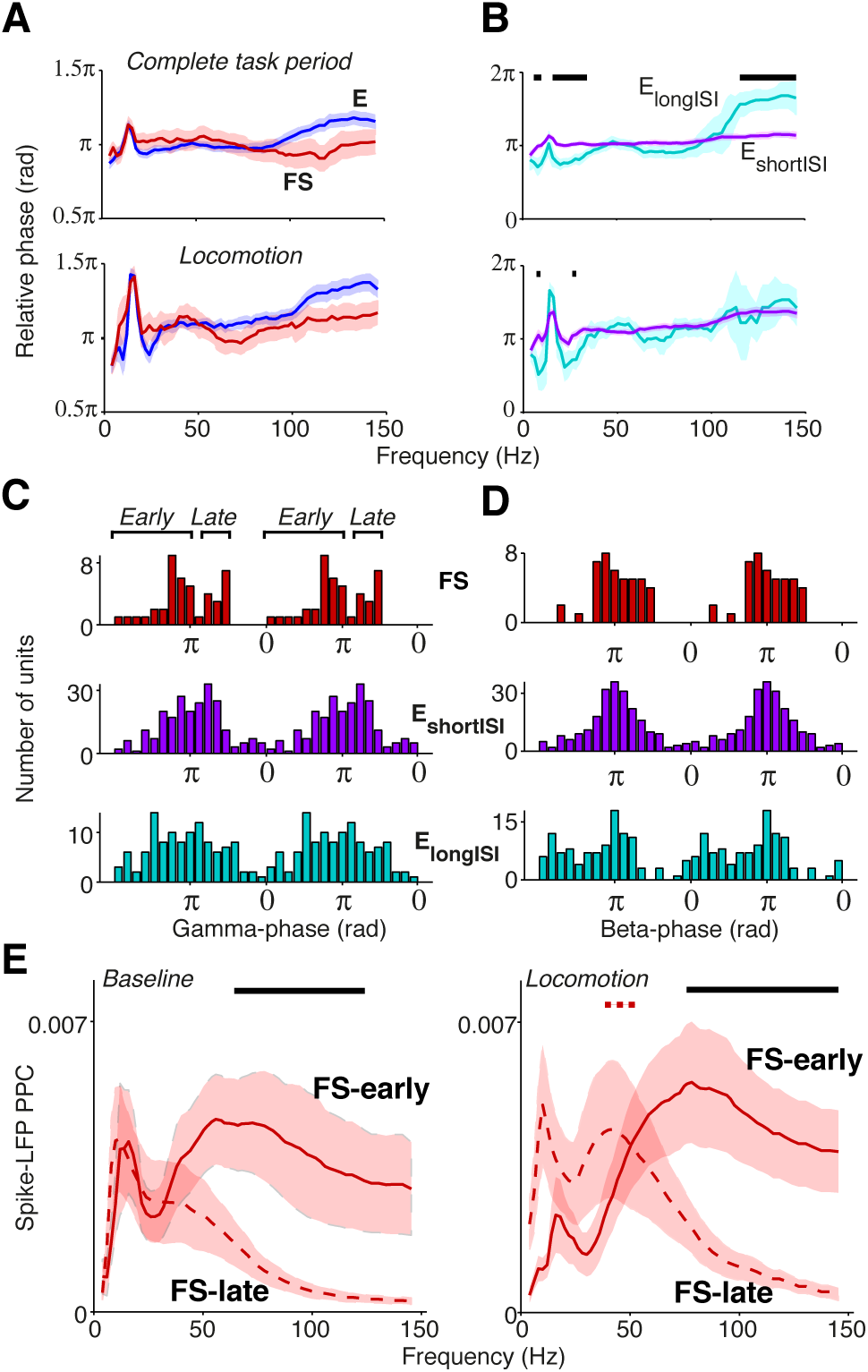
Phase relationships of distinct S1BF cell classes to S1BF LFP. **(A)** Mean spike-LFP phase for FS and E cells, separately for the complete task period and locomotion. At none of the frequencies did we find a significant difference in preferred phase of firing between FS and E cells. Shadings indicate 72% confidence intervals. **(B)** Same as **(A)**, but now for E_longISI_ and E_shortISI_ cells. **(A-B)** Black, horizontal bars indicate significant mean phase differences between the plotted cell classes (Circular ANOVA, p<0.05). **(C)** Histogram of preferred phases of firing in gamma cycle (taken at 66 Hz). On average, no significant phase difference between cell classes was detected. FS cells had a significant bimodal distribution of gamma phases (at all frequencies between 42 and 100 Hz, p<0.05, Hartigan’s dip test). **(D)** Phase histogram at 18 Hz for FS, E_longISI_ and E_shortISI_ cells. **(E)** Mean ± SEM of Spike-LFP PPC spectra for FS cells firing early and late in the gamma cycle, separately for baseline and locomotion. Black horizontal bars indicate significant differences between PPC measures for FS-early and FS-late cells. Dashed red horizontal bar indicates significant difference between baseline and locomotion for late-firing cells (p<0.05, multiple comparison corrected for number of frequencies (Korn et al., 2004).

Population gamma phase histograms revealed a more complicated picture of the temporal dynamics of firing in the gamma cycle (Figure 6C). The distribution of preferred gamma phases of FS cells was significantly bimodal (Figure 6C), with a ‘late’ group of cells (N=14) firing at a mean gamma phase of 249.7 ± 9.74° (mean ± 95% c.i. at 66 Hz), i.e. after the E cells, and an ‘early’ group of cells firing at a mean gamma phase of 136.7 ± 14.36°, i.e. before the E cells (bimodality significant for all frequencies between 42 and 100 Hz, p<0.05, Hartigan’s dip test, not significant for frequencies outside this range). This bimodality was neither observed for E cells (Figure 6C) nor for lower frequencies (Figure 6D).

We further investigated the firing properties of the FS-early and FS-late cells. We found that FS-early cells did not have significantly different firing rates (mean ± SEM = 12.6 ± 3.1 Hz) in comparison to FS-late cells (mean ± SEM = 8.9 ± 2.0 Hz, p = 0.24, Rank-Wilcoxon Test). Further, their LV values did not differ significantly (early: 0.86 ± 0.036; late: 0.82 ± 0.06, p = 0.4, Rank-Wilcoxon Test). This suggests that both FS cell classes are indeed GABAergic. We also found that nearly all FS-late cells were recorded from superficial/granular layers (L5/L6: 1; L4–5 transition: 0, L2–4: 13) whereas FS-early cells were relatively more evenly spread across layers (L5/6: 3, L4–5 transition: 9; L2–4: 16; p<0.05 for L4/5 transition and L2–4, randomization test).

Given their timing relative to excitatory cells, we hypothesized that the phase locking of FS-late cells to the gamma rhythm could depend on synchronous excitatory feedback, whereas the phase locking of FS-early cells should not. This makes two predictions: 1) the phase locking of FS-late cells should more closely resemble the phase locking of E cells, and 2) the phase locking of the FS-late cells should show a similar dependence on behavioral state as that of the excitatory cells (Figure 5A), whereas the locking of the FS-early cells should not. Indeed, we observed that FS-early cells were more strongly gamma phase locking than FS-late cells (Figure 6E) (p<0.05, MCC randomization test). Furthermore, just like the E cells (Figure 5A), the FS-late cells increased their gamma locking during locomotion while the FS-early cells did not (MCC randomization test, Figure 6E). In conclusion, our spike phase analysis reveals two subgroups of FS neurons that fire early versus late relative to excitatory neurons; the late-firing subgroup appears to follow excitatory cells in its dependence on behavioral state.

### Extracellular dataset: Inter-areal phase-synchronization

The analyses above show state and cell-type specific synchronization patterns within area S1BF, but do not yet elucidate how groups of S1BF neurons communicate with connected brain areas, as indicated by coherence in oscillatory activity. Therefore, our next question was how S1BF cells were phase locked to LFPs in the other recorded areas, namely the dorsal CA1 area of the hippocampus, perirhinal cortex (area 35/36), and area V1M. These areas have known mono- or disynaptic connections with S1BF (Aronoff et al., 2010; Naber et al., 1999; Paperna & Malach, 1991).

We first analyzed spike-LFP phase locking patterns, because the local origin of spikes is undisputed. For each of the four areas, we found spike-LFP gamma-synchronization that was enhanced during the locomotion period (Figure 7). The findings on S1BF gamma have been discussed in detail above. CA1 E (excitatory) cells were phase locked to CA1 LFPs especially at frequencies above 75 Hz (Figure 7), while CA1 FS (Fast Spiking) cells showed prominent spike-LFP phase locking at frequencies above 50 Hz, peaking around 100 Hz, and were more strongly phase locked in a broad gamma-band frequency range than E cells both during baseline and locomotion periods (Figure 7). Phase locking of E cells increased during locomotion (Figure 7), indicating that high-frequency locking did not reflect sharp-wave ripple activity (Csicsvari et al., 2003; Schomburg et al., 2014). Perirhinal cortex showed gamma phase locking to the local LFP in FS cells and E cells, with a relatively broad spectrum for E cells and more pronounced peak for FS cells (Figure 7). Gamma locking tended to increase with locomotion but this did not reach significance after multiple comparison correction (Figure 7). V1M cells were gamma locked around 50 Hz, which agrees well with previous reports in mice (Niell & Stryker, 2010; Vinck et al., 2015), carnivores and primates (Gray et al., 1989; Maldonado et al., 2000), and gamma locking was enhanced in the locomotion period (Figure 7). For each of the areas, local gamma-synchronization was also revealed using LFP-LFP coherence analysis (Figure 9). Thus, we conclude that significant gamma-band and supra-gamma synchronization could be detected in each of the four individual areas.

**Figure 7:**
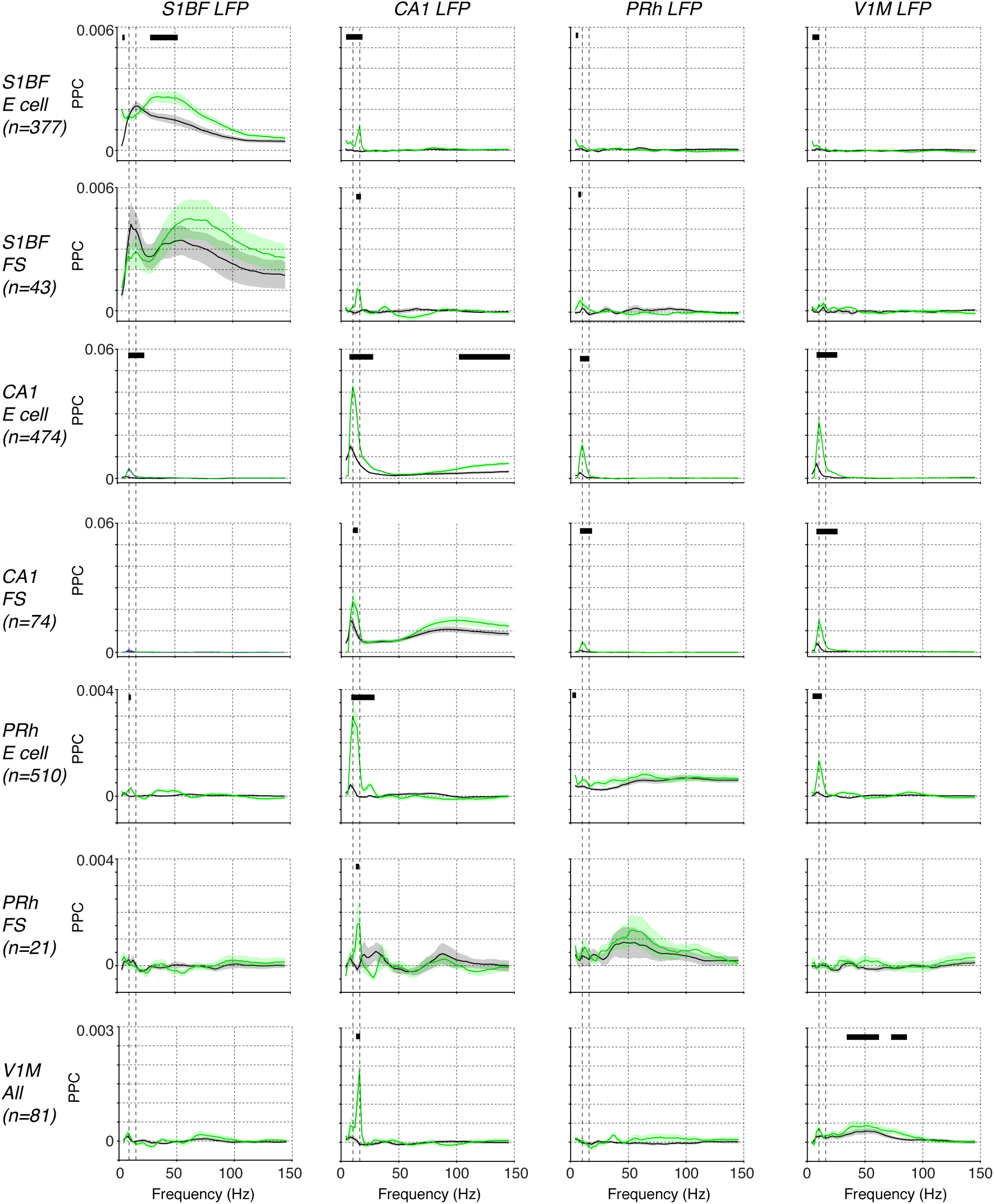
Phase locking among S1BF, CA1, V1M and perirhinal spikes and LFPs. Shown the mean ± SEM of spike-LFP PPC (Pairwise Phase Consistency) phase locking spectra, as a function of frequency (Hz). Displayed are locking spectra for S1BF FS and E cells, CA1 FS and E cells, PRh FS and E cells, and visual cortex cells. We pooled FS and E cells in visual cortex together because of the small sample size. Black horizontal lines indicate that two conditions held true: (1) there was a significant difference between baseline and locomotion (p<0.05), (2) the PPC values for the period with the highest locking values exceeded zero (p<0.05, multiple comparison corrected randomization test, Korn et al. (2004)). The second condition was used because a significant difference between baseline and locomotion would not have a meaningful interpretation if the PPC did not exceed zero in at least one condition. Vertical dashed lines indicate 8 and 14 Hz peaks.

In contrast to the occurrence of local gamma-synchronization in each of the four brain areas during both baseline and locomotion periods, we found a striking lack of inter-areal gamma synchronization among the four studied brain areas (Figure 7). Cells in S1BF were not significantly gamma locked to LFPs in the other areas. Vice versa, we did not observe significant gamma locking cells in the other areas to S1BF LFPs (Figure 7). The observation that there was no significant long-range gamma-synchronization also held true for other inter-areal combinations, and for the behavioral period in which gamma synchronization was strongest in each of the respective areas, i.e. the locomotion period (Figure 7). This negative result was not due to lack of statistical power, because we recorded hundreds of cells in each of the areas, and we found that PPC values in the gamma range were extremely small and therefore unlikely to be physiologically relevant (on the order of 10^−5^). LFP-LFP coherence patterns also did not reveal gamma-band peaks in the coherence spectrum during locomotion or baseline (Figure 7). We thus conclude that long-range interactions during the behaviors studied here do not rely on phase-coupling between gamma oscillators associated with each of the four areas, and that the gamma synchronization observed in each area does not measurably propagate in the present behavioral context. Apparently, even in a system with known anatomical connections, inter-areal gamma coherence can be lacking in the face of strong local gamma coherence in each individual area.

However, we did find evidence for state-dependent long-range low-frequency (<20 Hz) synchronization (Figure 7). During the baseline condition, we did not observe significant long-range low-frequency synchronization between S1BF cells and the other areas (Figure 7). However, during locomotion, we found that S1BF cells showed small but significant <20 Hz locking to the CA1 LFP, with a peak in the beta-band (14–16 Hz; Figure 7). In addition, we found that S1BF locking increased in delta frequencies to perirhinal and visual cortex. Moreover, during locomotion, cells in V1M locked to beta fluctuations in the CA1 LFP. In the same behavioral state, perirhinal cells showed an increase in theta and beta locking to the CA1 LFP. These increases in locking to area CA1 occurred in the absence of significant increases in local low-frequency synchronization in S1BF, V1M and perirhinal cortex (Figure 7). However, during locomotion, CA1 cells strongly increased theta phase locking to the local CA1 LFP. The two latter findings suggest that during locomotion, CA1 theta and beta fluctuations entrain spiking activity in the neocortex.

Coming back to inter-areal synchronization of S1BF, the low-frequency entrainment of S1BF cells to CA1 LFPs might be specific to a cell class, or it might be homogeneous across cell classes. To examine this, we analyzed the locking of the two classes of excitatory cells (E_shortISI_ and E_longISI_) and two classes of FS cells (i.e., early and late-firing) to CA1 LFPs. All S1BF cell classes were entrained by the CA1 theta/beta rhythm, with only small and non-significant differences between these classes at 14–16 Hz (Figure 8A). Furthermore we found clear phase preferences of S1BF cells relative to CA1 beta oscillations and we did not detect any significant difference in the preferred phase of spiking among the four cell classes (Figure 8B).

**Figure 8:**
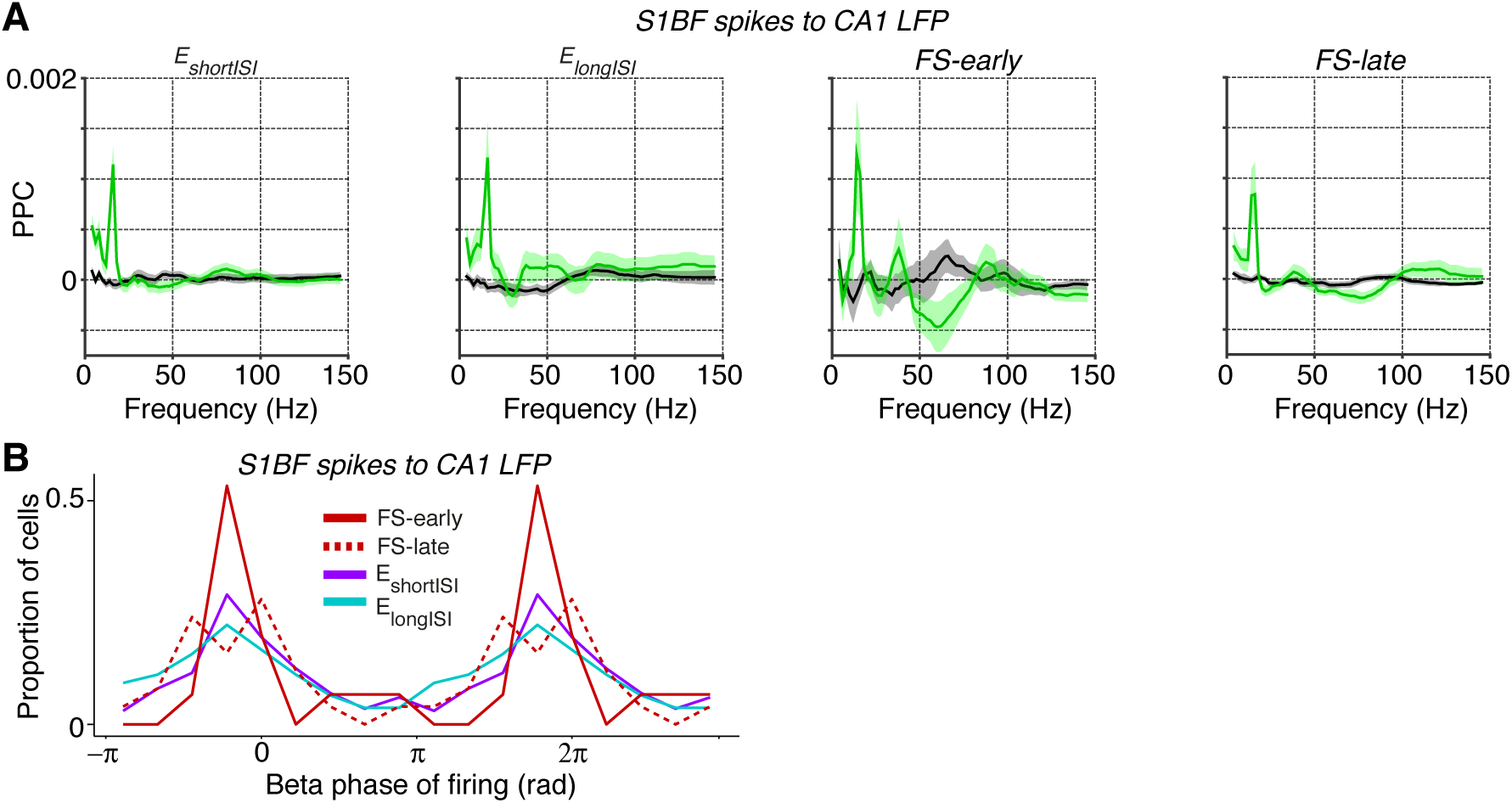
Spike-LFP phase locking between distinct S1BF cells and CA1 LFPs. **(A)** Entrainment of S1BF E_shortISI_, Eiongisi, FS-early and FS-late cells to CA1 LFPs, separately for locomotion and baseline. **(B)** Population histogram of preferred phases of firing for all FS-early, FS-late, E_shortISI_ and E_longISI_ cells to CA1 beta fluctuations (at 14 Hz, the peak in the phase locking spectrum). Y-axis indicates the proportion of cells. The beta cycle is shown twice for visualization purposes. No significant difference was observed in the preferred phases between the four cell classes (Circular ANOVA, p=0.47).

Our spike-LFP analysis also showed that CA1 cells were theta phase locked to S1BF, perirhinal cortex and V1M LFPs (Figure 7). In addition, we found strong LFP-LFP theta synchronization among S1BF, V1M and CA1 LFPs (Figure 9). However, our data and the data of Sirota et al. (2008) indicate that this likely reflects volume conduction of hippocampal currents to cortical EEG. The hippocampus generates high-amplitude theta-frequency potentials that slowly decay with distance due to the geometry of the hippocampus. Because multiple out-of-phase theta oscillators exist in the hippocampus, the volume conducted hippocampal theta currents can often be out of phase at nearby locations in the cortex (Sirota et al., 2008). The use of a local reference does therefore not fully eliminate the possibility that hippocampal theta currents contribute, even at a few millimeters distance. Furthermore, volume conduction of phase-heterogeneous hippocampal theta can cause LFP-LFP theta phase-synchronization between nearby electrodes to occur at a phase delay, leading to spurious peaks in WPLI spectra. Indeed, several aspects of our data suggest that CA1 cells phase lock to the volume conducted ‘shadow’ of hippocampal EEG dipoles: 1) While CA1 cells showed quite strong phase locking to theta fluctuations in S1BF and V1M LFPs, we did not observe a prominent theta peak in phase locking of S1BF and V1M cells to CA1 (with very low theta PPC values), nor did we detect theta peaks in the local spike-LFP phase locking spectrum in V1M and S1BF (Figure 7). 2) CA1 cells locked to cortical EEG depending on distance. They were only weakly locked to S1BF theta-band fluctuations, for which the (mostly superficial) recording sites lie furthest away from CA1, while they were strongly phase locked to theta fluctuations in V1M and perirhinal cortex, for which the recording sites lie closer to CA1 (Figure 7). This interpretation is consistent with the finding that especially V1M power spectra showed similar theta harmonics as the CA1 LFP power spectrum (Figure 9A), despite the absence of theta phase locking in V1M cells to either V1M LFPs or CA1 LFPs (Figure 7). (3) CA1 spike phase locking to neocortical LFPs occurred both during locomotion and baseline, while phase locking of neocortical cells to CA1 LFPs only occurred during locomotion (Figure 7).

**Figure 9:**
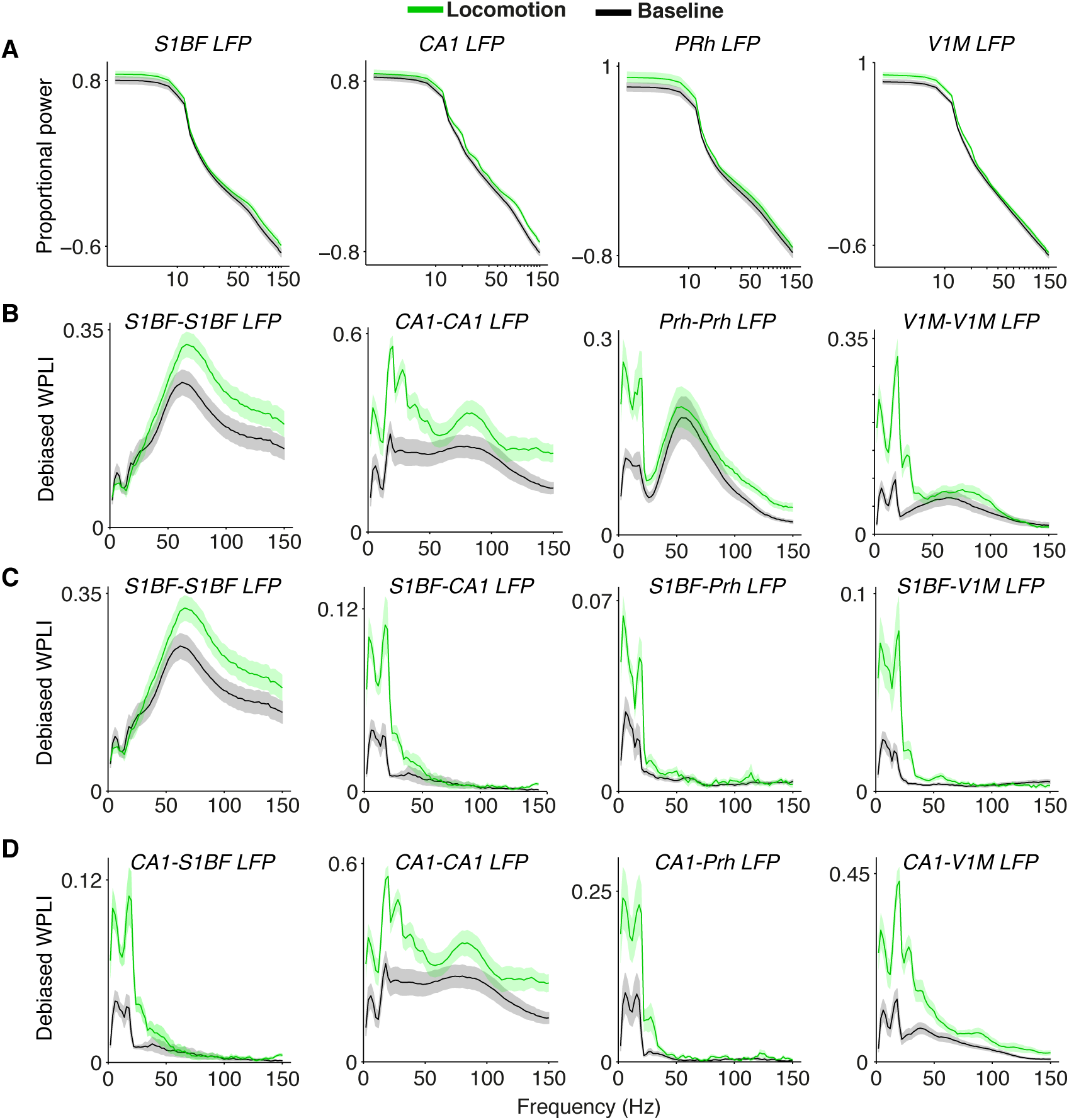
Inter and intra-areal phase-synchronization among S1BF, CA1, V1M and perirhinal cortex LFPs. **(A)** Normalized power spectra for the four areas, similar to Figure 1. **(B)** Phase-synchronization [debiased WPLI] between LFPs from the same areas, as in Figure 2. Shadings indicate SEMs across sessions. **(C**) Phase-synchronization between S1BF LFPs and LFPs from the other areas. First column is shown also in **(B)**. **(D)** Phase-synchronization between CA1 LFPs and LFPs from other areas. First and second column are also shown in **(B-C)**.

The likely influence of volume conduction of hippocampal currents on spike-LFP and WPLI connectivity metrics illustrates that the spatial origin of inter-areal interactions is not necessarily local and must always be carefully interpreted (Nolte et al., 2004; Schoffelen & Gross, 2014; Sirota et al., 2008; Vinck et al., 2011).

### Intracellular dataset: Modulation of gamma power in whole-cell recordings

We analyzed another dataset, first to validate our extracellular cell classification and, second, to examine whether a modulation of S1BF gamma-band synchronization by behavioral state can also be observed in the subthreshold membrane potential of inhibitory and excitatory neurons. Because the extracellular recordings did not yield data specifically on whisking activity, intracellular recordings in head-fixed mice were used in a complementary fashion to examine behavioral states in terms of whisking in free-air and active touch. If behaviorally modulated gamma activity can be confirmed, this would not only support the local expression of gamma activity in S1BF, but also indicate involvement of identified cellular phenotypes. However, because our analysis of membrane potential required that spikes be removed from the traces, it should be noted that no phase locking could be studied here.

First, our analysis of the data of Gentet et al. (2010, 2012) (Figure 10) showed that two clusters of cells can be reliably separated from each other, with one cluster consisting of SSt and PV cells, and another cluster consisting of non-fast spiking interneurons and excitatory cells. Both PV and SSt interneurons had, on average, higher firing rates than excitatory cells (p<0.001, Rank Wilcoxon test, Figure 10C). This analysis supports our interpretation of the extracellularly recorded FS cells (Figure 3) as containing two subgroups: PV and SSt cells.

**Figure 10:**
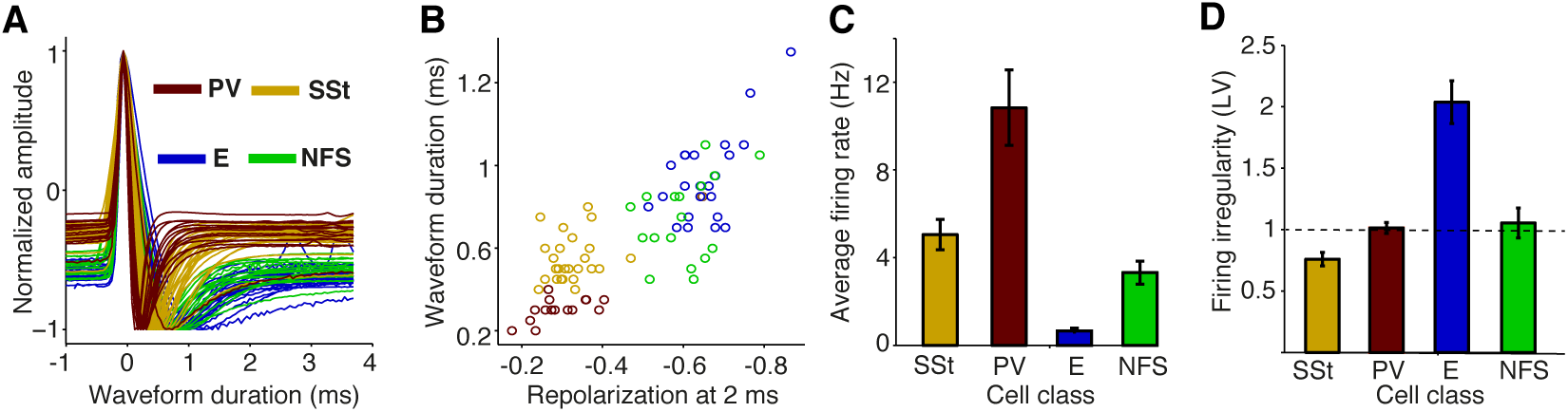
Action potential waveform and firing rate properties for identified cellular phenotypes in mouse S1BF. **(A)** AP (Action Potential) waveforms were obtained by whole-cell recordings from identified cellular phenotypes in awake mice. We took the 1st derivative of the AP waveforms to allow for a comparison with extracellular waveforms. Shown is the normalized amplitude of the 1st derivative as a function of time (ms) relative to peak. SSt: Somatostatin-positive; PV: Parvalbumin-positive; E: excitatory; NFS: non-fast spiking. **(B)** Normalized 1st derivative amplitude at 2 ms (repolarization) vs. peak-to-trough duration (ms). **(C)** Average firing rate (Hz) for different cell classes. **(D)** Mean ± SEM of firing irregularity (LV) for different cell classes.

Second, *V_m_* power spectra had an overall shape similar to the extracellular LFP recordings, with a dampening of the 1/*f^α^* trend below 10 Hz, and a 1/*f^α^* trend for higher frequencies. In contrast to the diverse effects that whisking has on firing rates of different cell types (in particular suppression and activation of SSt and NFS cells, respectively; Gentet et al. (2012)), it had quite uniform effects on the spectral characteristics of membrane potentials. Congruent with previous work (Gentet et al., 2010, 2012; Poulet et al., 2012; Zagha et al., 2013), we found that free-air whisking decreased low-frequency (1–5 Hz) fluctuations in the membrane potential of the various cell types relative to wake quiescence. However, free-air whisking significantly increased power in a broad range of frequencies in the 30 - 90 Hz range (p<0.001 for all cell types, Rank-Wilcoxon Test, N = 26, 23, 14 and 17 for SSt, PV, E and NFS cells), with spectral peaks between 30–50 Hz for SSt, excitatory and PV cells, and a 25–35 Hz peak for NFS cells (Figure 11A-B). Removing -2 to 10 ms of data around each AP for pyramidal cells led to the same conclusion, and for our Brownian noise control (see Material and Methods) no differences between free-air whisking and baseline periods were observed (Figure 11C-D). Notably, no significant differences were observed between cell types at 30–90 Hz (two-way ANOVA, n.s.), suggesting that the gamma increase was carried by common inputs to all of these superficial cells. We found a tendency for *V_m_* depolarizations and gamma increments to go hand in hand (Pearson’s R = 0.27, p<0.05, Figure 11E), indicating that cells expressing stronger gamma-rhythmic activity receive more depolarizing inputs. Moreover, active touch led to a further elevation of gamma-band power in comparison to free-air whisking (p<0.05, Rank-Wilcoxon test, N=8 ; Figure 11F-H), although the sample analyzed mostly consisted of SSt cells.

**Figure 11:**
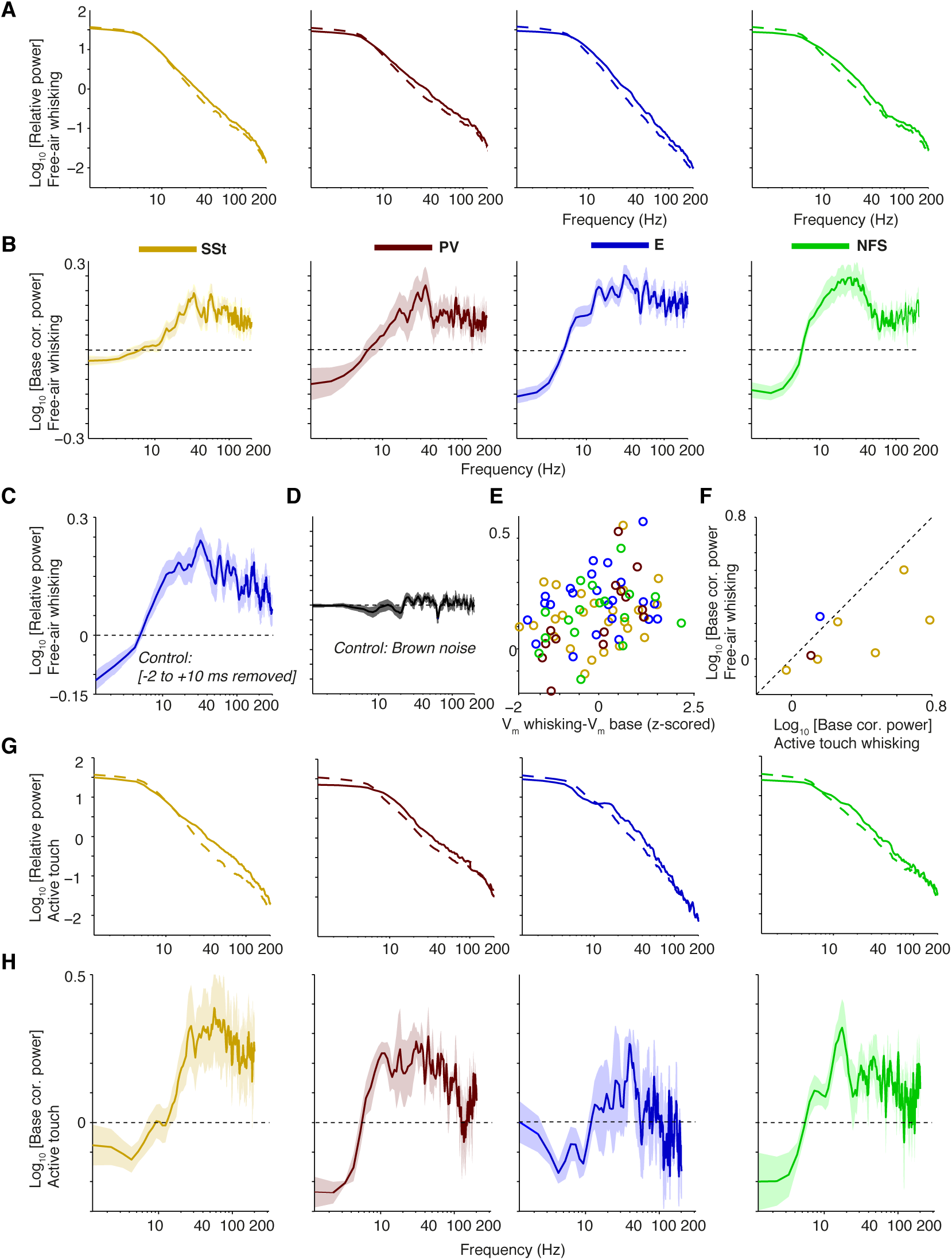
Gamma power increases in the *V_m_* of whole-cell recorded cell types during free-air whisking and active touch in S1BF of the mouse. **(A)** From left-to-right: relative power spectra (relative to sum of power across frequencies during baseline) for free-air whisking (solid) and baseline (dashed) for SSt interneurons, FS PV interneurons, excitatory, and NFS GABAergic interneurons. Shadings indicate SEMs. **(B)** As **(A)**, but now shown power change during free-air whisking relative to baseline (log_10_ transformed) **(C)** Same as **(A)**, third column, but now removing -2 to 10 ms around every spike. **(D)** Power change during free-air whisking relative to baseline (log_10_) for all cells together where original traces have been replaced by random Brownian 1 /*f^2^* noise. **(E)** Power change during free-air whisking relative to baseline (log_10_) as a function of the Z-scored difference in mean membrane potential difference between free-air whisking and baseline. **(F)** Power change during active touch whisking relative to baseline (log_10_) vs. power change during free-air whisking relative to baseline (log_10_). **(G-H)** As **(A-B)**, but now for active touch. **(H)** shown on same scale as **(B)**.

Thus, during active states, gamma-band power does not only increase in LFP signals, but also increases in the membrane potential of excitatory cells and multiple inhibitory interneuron classes. These findings complement the extracellular results in several ways, first showing that FS cells consist of two subclasses of interneurons (SSt and PV cells) and, second, that a state-dependent modulation of gamma activity is found in subthreshold membrane activity of both inhibitory and excitatory cell types.

## 4. Discussion

In this study we focused on better understanding how cells in neocortex are synchronized to oscillatory activity at various frequencies locally and in connected areas, and to what extent this depends on cell class and behavioral state. We examined this topic in S1BF, making simultaneous extracellular field and single-unit recordings from visual and perirhinal cortex as well as area CA1 in freely moving rats. In addition, we analyzed rhythmicity in whole-cell S1BF recordings in awake, head-fixed mice. Both gamma and beta rhythms entrained local S1BF spiking activity in a cell-type and state-dependent manner. In particular, we found differences between multiple S1BF inhibitory and excitatory cell classes in terms of (1) phase locking strength, (2) preferred phase of firing, and (3) behavioral state-dependence of locking (Figures 3–6). We did not find long-range gamma-synchronization among the four areas studied, even though gamma-band synchronization was detected in all four recorded areas individually. In contrast, we found inter-areal synchronization in the theta and beta ranges. In particular, S1BF cell firing was entrained by CA1 beta oscillations during locomotion.

### Gamma oscillations in S1BF

It is of note that while S1BF was the first brain area in which gamma oscillations were induced through the optogenetic activation of PV cells (Cardin et al., 2009; Siegle et al., 2014), there existed surprisingly little data indicating that, under naturalistic conditions, band-limited gamma oscillations occur in area S1BF. Few studies have shown modulation of LFP gamma-band power with whisking or vibrissae stimulation (Hamada et al., 1999; Jones & Barth, 1997; Zagha et al., 2013) or perceptual discrimination (Siegle et al., 2014). Studies including intracellular membrane potential data did not reveal signatures of gamma-synchronization (Poulet & Petersen, 2008), which is likely a consequence of a difference in analytical methodology. Moreover, none of the studies using extracellularly recorded signals (Hamada et al., 1999; Jones & Barth, 1997; O’connor et al., 2002; Siegle et al., 2014; Zagha et al., 2013) inquired whether raw LFP power or spike/LFP-LFP phase-synchronization spectra contain a peak in the gamma-frequency band, or whether enhancements in gamma-band power merely reflect broad-band or spiking-related increases in LFP power (Buzsaki et al., 2012; Buzsaki & Wang, 2012; Miller et al., 2009; Ray & Maunsell, 2011).

Might some of the S1BF gamma-rhythmicity that we report here be driven by the whisker kinetics that are caused by the rhythmic sweeping of the whisker or resonant whisker vibrations? Practically, it is very difficult to track the precise kinetics of the whiskers in freely moving rats: One needs a high resolution camera operating at high sampling rate, which can only focus on a very small region of space and image a single whisker for a limited amount of time, which likely requires whisker clipping. The analysis of this type of data is also far from trivial (Ritt et al., 2008; von Heimendahl et al., 2007; Wolfe et al., 2008). Even then, all the contributions from other whiskers (contextual modulation) might be missed, which likely is important given that gamma-synchronization is sensitive to contextual modulation in other areas (Gieselmann & Thiele, 2008). Thus, this approach is, at present, not practically feasible for the naturalistic behaviors studied here. The two major contributions to whisker kinetics are: (i) active whisker movements (e.g. Crochet & Petersen (2006); Wolfe et al. (2008)), and (ii) passive whisker vibrations through resonance (Lottem & Azouz, 2008; Ritt et al., 2008; Wolfe et al., 2008). Could this type of whisker kinetics explain the entrainment to LFP gamma oscillations?

As for the active whisker movements, whisker kinetics during free-air whisking are dominated by low frequencies, in particular a 8–10 Hz whisking cycle (Crochet & Petersen, 2006; Wolfe et al., 2008). Thus, it is unlikely that this alone is driving the gamma-synchronization observed. Furthermore, there might be a contribution to higher frequencies from resonant whisker vibrations arising when the rat is touching a texture or is free-air whisking (Ritt et al., 2008; Wolfe et al., 2008). Currently there is substantial debate as to how whisker kinetics encode different textures and contribute to perception and decision making (Lottem & Azouz, 2008; Ritt et al., 2008; von Heimendahl et al., 2007; Wolfe et al., 2008). Nevertheless, studies agree on a few points: (i) When, during active whisking, whiskers make contact with surface, they exhibit high-energy slip-stick events followed by transient ringing of the whisker at a characteristic resonant frequency (‘micromotions’) (Lottem & Azouz, 2008; Ritt et al., 2008; von Heimendahl et al., 2007; Wolfe et al., 2008); (ii) The resonant frequency depends on the whisker length, but generally ranges roughly between 100–300 Hz (Lottem & Azouz, 2008; Ritt et al., 2008; Siegle et al., 2014; Wolfe et al., 2008); (iii) Finally, the resonant frequency does not seem to be texture-specific, but rather the number and energy of slip-stick events is texture dependent (Lottem & Azouz, 2008; Ritt et al., 2008; Wolfe et al., 2008).

Given the stereotactical coordinates we used for implantation, our recordings were most likely made from around whisker C2/3 and D2/3, and they would extend across multiple barrel columns given the spatial extend of our recordings (our tetrode bundle width was 1 mm). B1–3, C1–3, D1–3 whiskers have prominent spectral peaks at frequencies higher than the classic gamma-range, i.e. between 100–300 Hz (Lottem & Azouz, 2008; Ritt et al., 2008; Wolfe et al., 2008). If whisker micromotions would be responsible for the observed gamma oscillations, then we would expect that: (i) gamma is extremely non-stationary, and only occurs during slip-stick events when whiskers make contact, with a characteristic dampened oscillation/ringing following slip-stick events; (ii) gamma would have its energy concentrated in the 100–300 Hz band. Rather, we observe that gamma has its energy concentrated in a lower-frequency band (50–70 Hz) and does not have the highly non-stationary character expected from the slip-stick events. Furthermore, we did not observe a difference in gamma power between locomotion trials with rough and smooth sandpaper on the walls. Nevertheless, it is to be expected that some of the spike-field phase locking at frequencies >100 Hz, which occurs esp. during the locomotion phase of the task, might be driven by neuronal activity triggered by whisker vibrations (Lottem & Azouz, 2008); future research is needed to investigate this.

Another possibility is that whisker touches evokes a gamma-rhythmic response in both the LFP and the spikes, and that this accounts for the observed phase locking. Evoked responses to self-initiated or passive whisker touches are extremely short-lived (Arabzadeh et al., 2005; Gentet et al., 2012; Lottem & Azouz, 2008) and show no evidence of an oscillatory pattern, however. Given that these responses can be roughly described as delta pulses convolved with an exponential or alpha-function, one expects them to have a broadband spectral profile with the energy focused in the low frequencies. There is also evidence that neurons mainly respond to the initial phase of whisker touch, rather than to the oscillatory ringing that occurs after the slip (Lottem & Azouz, 2008). These events would occur every 100–200 ms (whisker cycle duration), however inspection of the LFP and spike traces does not show evidence that gamma phase locking occurs in bouts of 100–200 ms. Furthermore, the sustained gamma-synchronicity observed in this study occurred throughout both the baseline and the locomotion phase, and had a sustained character, even though the rats engaged in quite different behaviors during these phases. This makes it unlikely that the gamma phase locking we observed was for a large part driven by evoked responses to whisker touch, however we cannot exclude this possibility with absolute certainty.

We found that the gamma-synchronization in S1BF covered a broad range of frequencies, which (as follows from the properties of the Fourier transform) means that the time-domain companions of the spike-field locking and field-field coherence measures tended to decorrelate rapidly over time (Figure 2D and Fig 4D). This means that the S1BF oscillation was - from the point of view of one single frequency - essentially short-lived. We found that the gamma oscillations in the other brain areas also had a broadband character. This broadband character of gamma is not atypical and has been reported for signals in visual cortex and hippocampus as well (Bragin et al., 1995; Burns et al., 2011; Schomburg et al., 2014; Vinck et al., 2013). For example, in visual cortex, gamma phase locking of FS cells extends from ≈30 Hz up to ≈120 Hz (and MUA-field locking is still non-zero at 120 Hz) (Fries et al., 2008; Vinck et al., 2013). In hippocampus CA1 and CA3, phase locking extends well above 150 Hz (Schomburg et al., 2014). In general, narrow-band phase locking can be seen when gamma is very strong in area V1, but it also tends to get more broadbandish under stimulus conditions when gamma is weaker (Gieselmann & Thiele, 2008). Thus, the phrases ’oscillation’ and ’rhythm’ used in this manuscript should not be understood in the sense that there is a narrow-band oscillator, but rather in the sense that the coherence spectra between the stochastic, wide-band field and spike signals show peaks in specific bands. The extent to which this imposes constraints on the potential function of gamma remains under active debate (Burns et al., 2011; Nikolić et al., 2013). Clearly, computational models of gamma and neuroscience theories should take actually reported - often weak - phase locking values and the broadband, often short-lived character of gamma into account.

Note that we were careful about never relating spikes to LFPs from the same electrode for the computation of phase locking. When there is bleed through of the spike to the field, there will be a sharp peak in the spike triggered average and a PPC spectrum that ramps up from about 30–50 Hz towards high frequencies, not a peak at gamma frequencies. We did not see this type of ramps when taking spikes from one electrode to the other electrode. Also, the inter electrode spacing was at least ≈150 micrometer, meaning that it was highly unlikely that spikes would have been measured on another tetrode, since the pick-up of spikes is spatially restricted to a few tens of micrometers (Gray et al., 1995). Finally, note that FS cells have narrower waveforms than E cells. This means that they contain less energy, and have their energy focused in a higher frequency band, since the spike can be described as an oscillation that has its energy mostly within the >300 Hz band. This means one would get less bleed-through with the FS cells at the gamma frequencies, and actually would predict less spurious inflation of the phase locking for FS than E cells. But we observed higher phase locking for FS cells than E cells.

We further note that there are interesting parallels between the gamma that we see in S1BF and the gamma that has been characterized in other cortical areas, such as area V4 of the macaque monkey: (i) the phase locking strengths of FS and E cells are comparable between S1BF and area V4 (during visual stimulation); (ii) phase-locking is broadband in both S1BF and area V4, with a spike-triggered average that decorrelates rapidly over time (Fries et al., 2001, 2008); (iii) the participation of E cells into the gamma-rhythms appears to be strongly dependent on network drive both in S1BF and area V4. There are also some notable differences, however, in particular in the distribution of FS phases (see below).

### Local vs. global beta and gamma synchronization

Until now only few studies have examined long-range gamma-synchronization between cortical sites using intracranial unit or field potential recordings, and those that have, provided mixed evidence as to whether gamma-synchronization forms a substrate for inter-areal communication (Bastos et al., 2015; Buschman & Miller, 2007; Engel et al., 1991; Gregoriou et al., 2009; Grothe et al., 2012; Schomburg et al., 2014; Sigurdsson et al., 2010; Sirota et al., 2008; van Kerkoerle et al., 2014; Von Stein & Sarnthein, 2000). In our dataset, gamma-synchronization was observed within all recorded areas, underscoring its ubiquitous occurrence in cortical circuits (Buzsáki, 2006; Fries, 2009). However, we found that S1BF gamma was not synchronized with the gamma rhythms in the other recorded areas. This lack of inter-areal synchronization occurred despite known anatomical connections between these areas and the previously reported occurrence of tactile responses in these other areas (Aronoff et al., 2010; Iurilli et al., 2012; Naber et al., 1999; Paperna & Malach, 1991; Pereira et al., 2007; Vasconcelos et al., 2011). Notably, we did not observe any significant gamma phase locking among the four studied areas, even though gamma-synchronization increased with locomotion in each individual area. Importantly, this conclusion was derived by making simultaneous unit and field recordings from multiple areas; as pointed out by Buzsáki & Schomburg (2015), field-field coherence patterns are often difficult to interpret because of common axonal projections (giving rise to phase shifted coherence) and because of volume conduction effects (Sirota et al., 2008). Taken together, our data argue against gamma-synchronization as a mechanism for bulk communication of tactile information during naturalistic behaviors, even though our data do not exclude the possibility that, under some task conditions, gamma coherence between these areas may selectively arise. Yet, the main point from this set of results is not that inter-areal gamma may never occur in the system studied; instead our results show that gamma synchronization can be lacking between interconnected brain structures in the face of gamma activity occurring in each area individually. This should prompt further research on the regulation of the coupling and uncoupling of interconnected oscillator systems (Buzsáki & Draguhn, 2004; Gielen et al., 2010; Kopell et al., 2000; Roberts et al., 2013; Steriade et al., 1993; Wang, 2010; Womelsdorf et al., 2014).

Inter-areal phase locking was most pronounced at lower frequencies, consistent with the idea that these frequencies are optimally suited for long-range coordination as they allow phase-coupling with longer interoscillator delays (Buzsáki, 2006; Kopell et al., 2000; Von Stein & Sarnthein, 2000). Sirota et al. (2008) showed that the envelope of neocortical gamma oscillations in rodents is modulated by hippocampal theta, and that cells in neocortex phase lock to CA1 theta oscillations. We found that during locomotion, but not during baseline, S1BF, V1M and perirhinal cortex activity was entrained by the CA1 EEG in the theta and beta band. This suggests that theta and beta rather than gamma phase locking coordinate the activity of hippocampal, parahippocampal and sensory areas during locomotion. Interestingly, both S1BF and V1M cell locking to CA1 LFPs occurred most prominently in the beta-frequency range. The beta-rhythmic entrainment of S1BF cells by CA1 LFP was homogeneous across cell classes, both in terms of phase locking strength and preferred phase of firing. This stands in contrast to the finding that local gamma synchronization in S1BF was strongly cell-type specific, in terms of (1) state-dependence of locking, (2) locking strength, and (3) phase of firing (Figures 3–8).

### Mechanisms of cortical gamma and beta oscillations

Models for gamma generation are commonly divided into ING and PING type of models (Bartos et al., 2007; Börgers & Kopell, 2005; Buzsaki & Wang, 2012; Eeckman & Freeman, 1990; Tiesinga & Sejnowski, 2009; Whittington et al., 1995; Wilson & Cowan, 1972). In ING models mutual I-I inhibition and gap junctions play a critical role in generating gamma (Bartos et al., 2007; Galarreta & Hestrin, 2001; Wang & Buzsaki, 1996; Whittington et al., 1995), without the requirement for synchronized E cell firing in the local circuitry. In PING models synchronous E-I activation plays a critical role in sustaining the rhythm (Börgers & Kopell, 2005; Csicsvari et al., 2003; Eeckman & Freeman, 1990; Wilson & Cowan, 1972). A strong prediction of the PING model is that within the gamma cycle, E cells fire first and trigger the firing of I cells, leading to a characteristic E over I phase lead (Börgers & Kopell, 2005; Wilson & Cowan, 1972). In ING models, I cells fire at the same gamma phase as E cells, or even at an earlier phase as I-I connections are relatively fast in comparison to I-E connections (Bartos et al., 2007; Whittington et al., 1995). Previous studies, using extracellular data from hippocampus, frontal and extrastriate cortex, have shown a phase lead of excitatory over FS cells (Csicsvari et al., 2003; Hasenstaub et al., 2005; Tukker et al., 2007; van Wingerden et al., 2010; Vinck et al., 2013). We however found that excitatory and FS cells fired, on average, at the same gamma phase. We might have merely failed to detect a phase difference, however the 95% confidence intervals of FS and excitatory cells’ phase distributions indicated a maximum delay of about 1 ms. At first glance, this finding appears inconsistent with the PING model, as the latter predicts a phase lead of excitatory over inhibitory cells. However, our data shows that FS cells do not form a homogeneous group, but consist of early-firing and late-firing cells. It remains unclear what accounts for the strong phase locking of FS-early cells. Given the weak synchrony of the E cells, one might hypothesize that the gamma-rhythmicity of the FS-early cells is established through mutual inhibition (an ING type of mechanism). The FS-early cells would then entrain the network through rhythmic inhibition, and the FS-late cells would inherit the gamma rhythm from the excitatory cells through E-I projections, causing a characteristic phase delay (Figure 12). The whole-cell recordings analyzed here indicate that the FS cell group should encompass both PV and SSt cells. Because PV basket cells have strong mutual inhibition and gap junction connections (Bartos et al., 2007), it makes them an ideal candidate to generate robust gamma oscillations via I-I connections in the absence of synchronous E activation (Bartos et al., 2007; Buzsaki & Wang, 2012); these cells would be the strongly gamma-locked, early firing cells. SSt cells do not possess strong mutual inhibitory connections, and do not receive a substantial projection from the PV basket cells (Pfeffer et al., 2013). Hence, they may rely exclusively on synchronous E cell activation to participate in the gamma rhythm; these cells would be the weakly gamma-locked, late firing cells. In turn, their activity might inhibit excitatory and FS basket cells late in the gamma cycle (Pfeffer et al., 2013).

**Figure 12:**
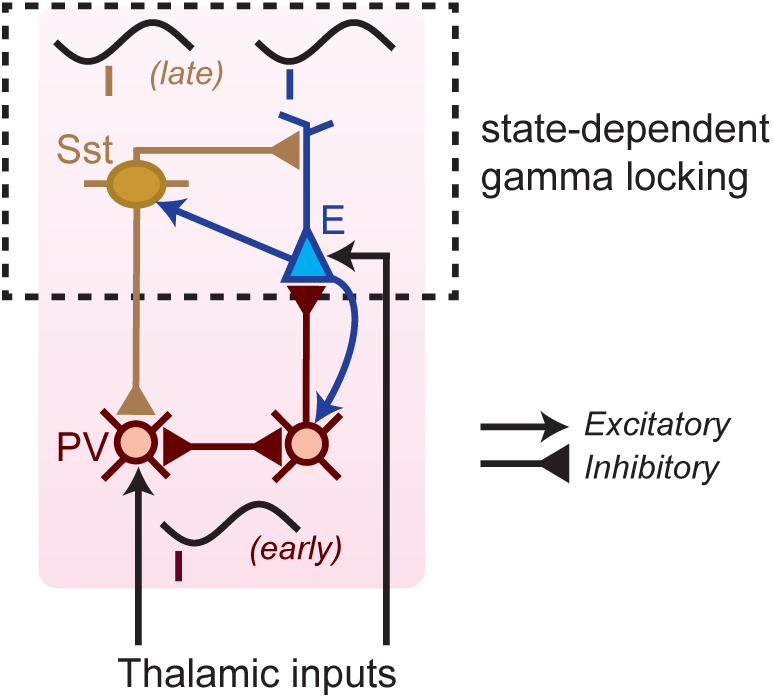
Hybrid model of S1BF gamma generation. A class of FS interneurons (red; presumably Parvalbumin-positive cells) has strong mutual inhibitory connections, and does not have a substantial projection to other FS interneurons (Pfeffer et al., 2013). These FS cells receive both excitatory drive from thalamic projections and from local recurrent excitatory inputs. Due to the mutual inhibitory connections, the FS cell network starts firing gamma synchronously and also entrains the excitatory cells. The excitatory cells fire at a slightly later phase because the inhibitory synaptic potentials from I→E connections are slower than the I→I connections (Bartos et al., 2007; Wang & Buzsaki, 1996). Another group of FS cells (yellow, presumably Somatostatin-positive cells) relies on the local excitatory cells to be entrained to the gamma rhythm, does not receive a substantial thalamic drive, and does not exhibit strong gamma phase locking. These cells fire late in the gamma cycle, few milliseconds after the excitatory cells, and might help to shut down the activity of both the early firing FS cells and the excitatory cells (Pfeffer et al., 2013). Early-firing FS cells show little state-dependence, whereas excitatory and late-firing FS cells show strong state dependence.

Our data also shed light on the generation of beta oscillations. We found that beta spike-LFP phase locking was particularly prominent in S1BF FS cells and in sparsely firing, bursting excitatory cells (Figure 5). Theoretical work has indicated that irregularly bursting cells indeed play a role in the generation of cortical beta oscillations (Roopun et al., 2008; Womelsdorf et al., 2014), and previous work had already shown that beta spike-field coherence is most prominent in deep layers of visual cortex (Buffalo et al., 2011), in which irregularly bursty cells tend to reside (McCormick et al., 1985). We found that S1BF beta-synchronization was found to be associated with deactivations (Figure 5), which is consistent with previous reports on increased beta-band synchronization in the period preceding motor execution (Donoghue et al., 1998; Murthy & Fetz, 1996). However, whereas local S1BF beta-synchronization was associated with deactivations, with the beta peak attenuating in the phase locking spectrum of both FS and E cells during locomotion, we found that during locomotion S1BF cells increased their beta phase locking to CA1 (Figure 7).

Recent work has highlighted the strong dependence of cortical activity on multiple awake behavioral states, such as quiescence, whisking, locomotion and arousal (Crochet & Petersen, 2006; Gentet et al., 2012; McCormick et al., 2015; Niell & Stryker, 2010; Vinck et al., 2015; Zagha et al., 2013; **?**). In agreement with these previous studies, our data indicate that S1BF gamma-synchronization is a signature of the active, information-processing state, as we found that gamma was enhanced both during locomotion (in terms of LFP power) and during whisking and active touch (in terms of *V_m_* power), was associated with firing rate increases, and was strongly quenched during slow wave sleep. Interestingly, we found the state-dependence of gamma-band synchronization to be cell-type specific. Late-firing FS cells had a strong S1BF state dependence (Figure 6), which might derive from the inputs of the excitatory cells whose gamma-rhythmic firing itself was state-dependent. This increase in E and FS-late gamma locking co-occurred with an increase in LFP gamma-band power and LFP-LFP phase synchronization. On the other hand, we found that the gamma phase locking of FS-early cells did not depend on behavioral state, indicating that it is, to some extent, robust against fluctuations in synchronization of the excitatory cells. This parallels the recent finding that in area V4 of the macaque, FS cells are gamma phase locking with similar strength in the stimulus and prestimulus period, whereas E cells are gamma phase locking only in the stimulus period (Vinck et al., 2013); the mechanisms underlying this differential state-dependence remain unknown.

## Acknowledgements

The authors would like to acknowledge the software tools provided by Dr. Kenneth Harris (Imperial College London, UK) for the use of KlustaKwik, and by Dr. A. David Redish (University of Minnesota, Minneapolis, MN) for the use of MClust. We thank Dr. Pascal Fries, Dr. Thilo Womelsdorf and Dr. Quentin Perrenoud for helpful comments on this work. We are grateful for the support provided by the technology center of the University of Amsterdam, in particular Gerrit Hardeman, Eric Hennes, Harry Beukers, Ron Manuputy, Matthijs Bakker, Ed de Water, and many others. We are grateful to Carl Petersen for allowing us to analyze data gathered in his lab. This work was supported by the Netherlands Organization for Scientific Research-VICI Grant 918.46.609 (to CMAP) and the EU FP7-ICT grant 270108 (to CMAP). MV, JJB and CMAP designed and planned the study. MV and JJB performed the quad-drive tetrode recordings, with support from LAD, JCJ, KTO and GAK. MV and JJB analyzed the extracellular tetrode data. LG performed whole-cell recordings, and MV analyzed this data. MV, JJB and CP wrote the paper, in collaboration with the other authors.

